# Live calcium imaging of *Aedes aegypti* neuronal tissues reveals differential importance of chemosensory systems for life-history-specific foraging strategies

**DOI:** 10.1101/345389

**Authors:** Michelle Bui, Jennifer Shyong, Eleanor K. Lutz, Ting Yang, Ming Li, Kenneth Truong, Ryan Arvidson, Anna Buchman, Jeffrey A. Riffell, Omar S. Akbari

**Author notes:** M.B., J.S, and E.K.L contributed equally to this work. To whom all correspondence should be addressed: Omar S. Akbari, Section of Cell and Developmental Biology, University of California, San Diego, La Jolla, CA 92093, USA, Jeffrey A. Riffell, Department of Biology, University of Washington, Seattle, WA 98195, USA.

## Abstract

*Aedes aegypti* have a wide variety of sensory pathways that have supported success as a species as well as a highly competent vector of numerous debilitating infectious pathogens. Investigations into mosquito sensory systems and their effects on behavior are valuable resources for the advancement of mosquito control strategies. Numerous studies have elucidated key aspects of mosquito sensory systems, however there remains critical gaps within the field. In particular, compared to that of the adult form, there has been a lack of studies directed towards the immature life stages. Additionally, although numerous studies have pinpointed specific sensory receptors as well as relevant response behaviors, there has been a lack of studies able to monitor both concurrently. To begin filling aforementioned gaps, here we engineered *Ae. aegypti* to ubiquitously express a genetically encoded calcium indicator, GCaMP6s. Using this strain, combined with advanced confocal microscopy, we were able to simultaneously measure live stimulus-evoked calcium responses in both neuronal and muscle cells with a wide spatial range and resolution. Moreover, by coupling *in vivo* calcium imaging with behavioral assays we were able to gain functional insights into how stimulus-evoked neural and muscle activities are represented, modulated, and transformed in mosquito larvae enabling us to elucidate mosquito sensorimotor properties important for life-history-specific foraging strategies.

**Significance Statement:** Understanding mosquito sensory systems and resulting behavior has been a major factor in the advancement of mosquito control innovations. *Aedes aegypti* larvae offer an effective life stage for further elucidating information on mosquito sensory systems. Due to their relatively simplified nervous system, mosquito larvae are ideal for studying neural signal transduction, coding, and behavior. Moreover, a better understanding of the larval sensory system may enable the development of novel control methodologies able to target mosquitoes before they reach a vector-competent stage. Here we engineer *Ae. aegypti* to ubiquitously express a genetically encoded calcium indicator, GCaMP6s and use this tool to observe links between sensorimotor responses and behavior by exploiting live calcium imaging as well as live tracking based behavioral assays.

## Introduction

The yellow fever mosquito, *Aedes aegypti*, is a global vector of numerous debilitating arboviruses including Chikungunya, Dengue, Yellow Fever, and Zika [1]. Due to its ability to transmit copious pathogens, adaptability to diverse climates, flexibility in oviposition sites, and desiccation-tolerant eggs, *Ae. aegypti* are significant worldwide epidemiological burdens, leading to hundreds of millions of infections annually resulting in over 50,000 deaths [2–5]. To decrease the imposed global burden, many vector control methodologies have been developed and implemented, including a number of innovative genetic-based technologies such as the release of insects carrying dominant lethal (RIDL)[6], and the infection and introduction of mosquitoes harboring the intracellular bacterium, *Wolbachia* either spread into populations to reduce viral transmission [7,8], or used for population suppression through *Wolbachia* induced cytoplasmic incompatibility [9]. Moreover, there are also a number of innovative “gene drive” based technologies that are rapidly being developed in *Ae. aegypti* with the hope of making an impact in the future [10–12]. Nonetheless, the most prevalent form of mosquito control used in the field today is the traditional use of chemical insecticides [13]. Although insecticides can have an impact on mosquito populations, due to their high costs, requirements for continuous application, and rapid susceptibility to resistance [14], they are not sustainable long-term solutions. Therefore, significant efforts are necessary to discern the underlying molecular, genetic and physiological mechanisms important for arboviral vector competence with the overall aim of developing additional novel, insecticide-free methods to disrupt viral disease cycles [15].

At both the larval and adult stages, mosquito sensory systems play pivotal roles in mediating a variety of behaviors, including locating food resources, habitat selection, and predator avoidance (Reviewed in [16–18]). As such, sensory systems provide attractive targets for suppressing vector behaviors at both the larval and adult stages. Over the years there have been numerous studies on adult mosquito sensory systems that have greatly advanced the field, such as the discovery of key olfactory and gustatory receptors [19,20] as well as behavioral responses to host cues [21–23]. However there remains critical gaps in understanding the direct relationships between sensorimotor and behavioral responses, specifically important for behaviors linked to vector competence such as host seeking and chemical avoidance. Additionally, only a handful of studies have focused on larval chemosensory systems resulting in significant gaps in a holistic understanding of mosquito sensory systems [24]. For example, olfaction is important for detecting long-range host cues in adult mosquitos. However in an aquatic environment, either gustation or olfaction could detect long-range food indicators [25]. Food scarcity is an important ecological constraint on mosquito larvae [26], but little is known about the chemosensory mechanism of foraging in larval mosquitoes. Given the relative simplicity of the larval nervous system, understanding chemosensory signal transduction, coding, and behavior in larvae could lead to novel control interventions and enable a more holistic understanding of mosquito behavior in areas such as food seeking and chemotaxis [17,24].

Notwithstanding, as of recently, we have lacked effective genetic tools to study mosquito larval sensory systems as they process environmental information. Current tools used in mosquitos to monitor neural activity include extracellular recording from sensilla and antennal lobe cells [27], as well as using synthetic calcium-sensitive dyes (e.g., FURA-2) *in vivo* or in heterologous systems [28,29]. To overcome the challenges of these existing approaches, here we have engineered *Ae. aegypti* to express a Genetically Encoded Calcium Indicator (GECI), termed GCaMP6s. GCaMP6s enables imaging of sensory-evoked calcium transients through changes in relative fluorescence [30]. Using this tool we were able to gain the unprecedented ability to concurrently measure *in vivo* sensory responses and motor responses with high spatial and temporal resolution in regions of neuropil and muscles of live responding mosquitos. This enabled us to gain functional insights into the importance of chemosensory channels in mediating behavior (e.g. foraging) by simply activating distinct olfactory and gustatory channels and measuring larval neural responses to diverse chemosensory stimuli in various genetic backgrounds such as those harboring mutations in important olfactory and gustatory receptors [19,20]. Taken together, our results link the sensory processing of specific stimuli to behavior responses of freely swimming larvae, thereby gaining a deeper functional understanding of mosquito multisensory integration.

## Results

### Development of an optogenetic-reporter of neuronal activity in *Ae. aegypti*

To visualize live calcium activity within *Ae. aegypti*, we engineered a transgenic *Ae. aegypti* strain harboring genomic sources of a genetically-encoded calcium indicator, GCaMP6s [30]. To express GCaMP6s, we utilized the *polyubiquitin* promoter (AAEL003877, henceforth *PUb*), chosen for its generally high expression during nearly all developmental life stages and tissues as shown by previous promoter characterization experiments and developmental transcriptional profiling (Figure 1A) [31,32]. We inserted the *PUb* promoter upstream of the coding sequence for GCaMP6s within a randomly inserting *piggybac* transposable element. Downstream to the *PUb* promoter driven GCaMP6s, we included an OpIE-2 promoter driving dsRed expression to serve as a robust transgenesis marker (Figure 1B). The engineered *piggybac* transgene was injected into the germ cells of preblastoderm stage *Ae. aegypti* embryos (0-1hr old). Transgenic G1 mosquitoes harboring the transgene were readily identified by a bright expression of OpIE-2 driven dsRed in the abdomen, in addition to a robust calcium signaled activation of GCaMP6s in muscle and neural cells (Figure 1C). To ensure that this transgenic line represented a single chromosomal insertion, we backcrossed isolated individuals for four generations to our wild-type (+/+) strain and measured Mendelian transmission ratios each generation and observed 50% of offspring inheriting the transgene, indicating that this strain represents a single chromosomal insertion. To determine its genomic insertion location, we used inverse PCR and found the location of insertion to be on the 2nd chromosome with flanking 5’ and 3’ *piggyBac* regions positioned at AaegL5.0 reference ( genomic loci 285,175,805-285,176,289 and 285,175,275-285,175,803, respectively.

**Figure 1.**
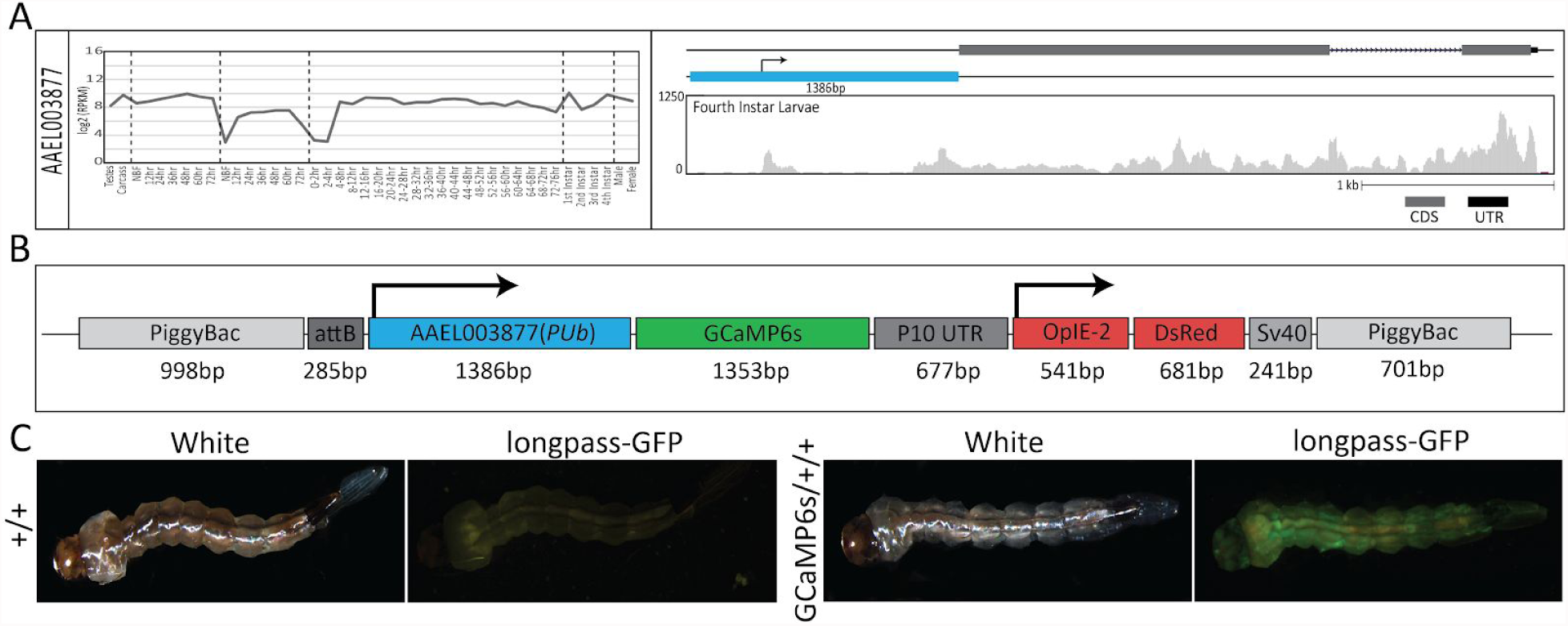
RNAseq expression, schematic representation of the GCaMP6s construct and larval fluorescence. Log_2_ (RPKM) expression values for the promoter, AAEL003877 (PUb) was plotted across development. Samples include, from left to right: testes; male carcasses (lacking testes); carcasses of females prior to blood feeding (NBF); female carcasses 12 hr, 24 hr, 36 hr, 48 hr, and 72 hrs post blood meal; ovaries from NBF females and at 12 hr, 24 hr, 36 hr, 48 hr, and 72 hr post ecdysis; embryos from 0–2 hrs through 72–76 hrs; whole larvae from 1st instar, 2nd instar, 3rd instar and 4th instar; male pupae; and female pupae. A genome browser snapshot was with the Y axis showing expression level based on raw read counts of fourth instar larvae (A). A schematic representation of the piggybac-mediated GCaMP6s construct. GCaMP6s is driven by AAEL003877(PUb)(blue) while dsRed by OpIE-2, the latter serving as a transgenic marker (B). Larval bright field images (*left*) and corresponding fluorescent images (*right*) show robust GFP transients throughout the whole body and DsRed fluorescence in the abdomen. +/+ represents wild-type larva. GCaMP6s/+/+ represents transgenic GCaMP6s larva (C).

### Temporal and Spatial Odor-Evoked GCaMP6s Responses in Adults

To assess GCaMP6s functionality in a wild-type genetic background (termed GCaMP6s/+/+ from hereon), and to visualize sensorimotor activity elicited by specific sensory channels, we initially recorded and quantified calcium transients in adult mosquitoes that were stimulated with CO_2_. Distinct regions-of-interest (ROIs) were imaged across various sensory organs of adult mosquitoes using laser-scanning confocal microscopy. Calcium-evoked changes in fluorescence varied between sensory organs tested. For example, the tip of the maxillary palp displayed significant changes in fluorescence intensity across 4 replicates presumably related to the location of olfactory sensory neurons (OSNs) within capitate peg sensilla on the maxillary palp [20] (meanΔF/F0 is 1.24 ± 0.13, p-value = 0.0323) (results from one replicate shown in Supplemental Figure 1A, Supplemental Video 1). While in the adult antennal flagellum, changes in fluorescence were recorded in the nodes between antennal segments (meanΔF/F0 is 0.04 ± 0.31, p-value = 0.8240 and 0.01 ± 0.12, p-value = 0.9469, for ROI 1&2 respectively) and the internodes (meanΔF/F0 is −0.08 ± 0.07, p-value = 0.1467), although neither were highly significant when comparing across 6 replicates (results from one replicate shown in Supplemental Figure 1B, Supplemental Video 2). Additionally, when observing clusters of ommatidia within the adult eyes, changes in calcium signaled GCaMP6s activation were highly stochastic across 4 replicates presumably due continuous optical responses (meanΔF/F0 is 0.02 ± 0.01, p-value = 0.4700; −0.15 ± 0.20, p-value = 0.4398; 0.02 ± 0.04, p-value = 0.4949, for ROIs 1-3 respectively) (results from one replicate shown in Supplemental Figure 1C, Supplemental Video 3). Although these results indicated that many regions (e.g. adult antennal flagellum and adult ommatidia) did not demonstrate significant odor-evoked responses across multiple replicates, accurate detection of subcuticular fluorescence was hindered by the adult’s thick cuticle and dense setation.

### Relative Odor-Evoked GCaMP6s Responses in Preadult Larval Stages

Compared to adults, 2nd instar larvae have relatively simplified neuroanatomical systems and a transparent cuticle making them more suited for detecting subcutaneous changes in fluorescence intensity reported by GCaMP6s. These factors coupled with the limited knowledge regarding mosquito larval sensory responses motivated us to simultaneously image muscle and sensory calcium-evoked responses with the GCaMP6s/+/+ strain. Using either a 5X or 10X objective permitted us to record fluorescence in the whole body, or just the head capsule, respectively. Results from these experiments revealed significant changes in calcium transients within the longitudinal muscles within the 2nd abdominal segment across 4 replicates (meanΔF/F0 is 3.45 ± 0.57, p-value = 0.0011) (results from one replicate shown in Supplemental Figure 1D, Supplemental Video 4) in the body in addition to the lateral retractors (mean ΔF/F0 is 2.04 ± 0.77, p-value = 0.004497), the deuterocerebrum (DE) across (mean ΔF/F0 is 0.45 ± 0.14, p-value = 0.002150) and medial retractors (mean ΔF/F0 is 2.57± 01.27, p-value = 0.013050) in the head across 5 replicates (results from one replicate shown in Supplemental Figure 1E, Supplemental Video 5). Taken together, these results indicate that GCaMP6s can be used to effectively visualize sensorimotor activity in neural and muscle tissues of live mosquito larvae.

### Calcium Imaging of Odor-evoked Responses in the Larval brain in Response to Olfactory Stimuli

To gain a more comprehensive understanding of the links between stimulus-evoked calcium responses in the brain and muscles of the larval head, a novel, minimally invasive, tethered-swimming assay was developed. This assay consisted of adhering the dorsal side of the larval head to a chambered cover glass, thereby immobilizing the head, while the larva was submerged in enough water to enable constant imaging of calcium transients within the head capsule while the larval tail could freely swim and its siphon could obtain oxygen (Figure 2A). Importantly, the larval head capsule is strikingly transparent requiring no surgical removal of cuticle thus enabling the larva to survive for extended periods (up to 48 h) enabling multiple recordings on the same individual. Responses in the DE and lateral abductors were analyzed as representatives of brain and muscle responses, respectively with at least 3 biological replicates per stimuli. Stimuli tested included chemicals previously shown to be relevant to adult mosquitoes such as a known repellent (VUAA1) [33], attractants (1-octen-3-ol, ethyl acetate and lactic acid) [34–37], a known exciter of multiple groove-peg ORNs (Butylamine) [25], and other behaviorally relevant compounds (sucrose, lobeline, glutamate, fish food) [38,39]. By stimulating GCaMP6s/+/+ larvae to this panel of chemicals we found that there were significant calcium responses in the DE to several stimuli including 1-octen-3-ol (max ΔF/F0 is 6.09 ± 3.85, p-value = 0.0145), butylamine (max ΔF/F0 is 3.29 ± 3.05, p-value = 0.0092), ethyl acetate (max ΔF/F0 is 3.14 ± 2.61, p-value = 0.0077), lobeline (max ΔF/F0 is 2.57±2.17, p-value = 0.0224), lactic acid (max ΔF/F0 is 2.31 ± 1.72, p-value = 0.0366), and VUAA1 (max ΔF/F0 is 2.12 ± 1.66, p-value = 0.0458) while sucrose (max ΔF/F0 is 2.01 ± 2.70, p-value = 0.1053), glutamate (max ΔF/F0 is 0.79 ± 0.61, p-value = 0.1632), and fish food extract (max ΔF/F0 is 0.62 ± 1.02, p-value = 0.6101) did not display significant changes in fluorescence intensity when compared to responses evoked by water (Figure 2D,2E, Supplementary Figure 2,3A). Interestingly, 1-octen-3-ol, a known mosquito adult attractant produced by microbes [40], displayed the greatest calcium response, followed by butylamine and ethyl acetate, with the former previously documented to induce a response in activated grooved-peg ORNs in *Anopheles gambiae*, *Anopheles quadriannulatus*, and *Culex quinquefasciatus*. Moreover, when observing muscle responses to the same stimuli we observed significant calcium increases in 5 of the 6 stimuli that also displayed significant responses in the DE. These stimuli included 1-octen-3-ol (max ΔF/F0 is 6.38 ± 1.50, p-value = 2.91e-05), butylamine (max ΔF/F0 is 6.11 ± 4.04, p-value = 4.11e-05), Ethyl Acetate (max ΔF/F0 is 3.10 ± 2.88, p-value = 0.00096), lobeline (max ΔF/F0 is 2.41 ± 2.45, p-value = 0.00408), and VUAA1 (max ΔF/F0 is 3.16 ± 2.16, p-value = 0.00437). Similar to the DE, the muscles did not exhibit significant responses to sucrose (max ΔF/F0 is 1.04 ± 1.64, p-value = 0.07634), glutamate (max ΔF/F0 is 0.68± 1.07, p-value = 0.1453), or fish food (max ΔF/F0 is 0.59 ± 0.45, p-value = 0.1534) when compared to responses evoked by water. Contrary to results from the DE, responses by muscles to lactic acid (max ΔF/F0 is 1.53 ± 1.88, p-value = 0.07509) were not significant. Lastly, the universal expression of GCaMP provided an opportunity for temporal comparison between brain and muscular responses When analyzing the latency in response between the DE and muscles (Figure 2 B,C), 1-octen-3-ol elicited a significant latency of 2.24 ± 3.19 sec (p-value = 0.003036), while butylamine elicited a latency of 0.48 ± 1.11 sec (p-value = 0.2867)(Figure 2F). Furthermore, a persistence in response to 1-octen-3-ol was seen in the DE but not in the muscle. This contrasted with the response to a majority of the other stimuli including Butylamine where fluorescent expression in the muscle matched tht of the DE (Supplementary Figure 7).

**Figure 2.**
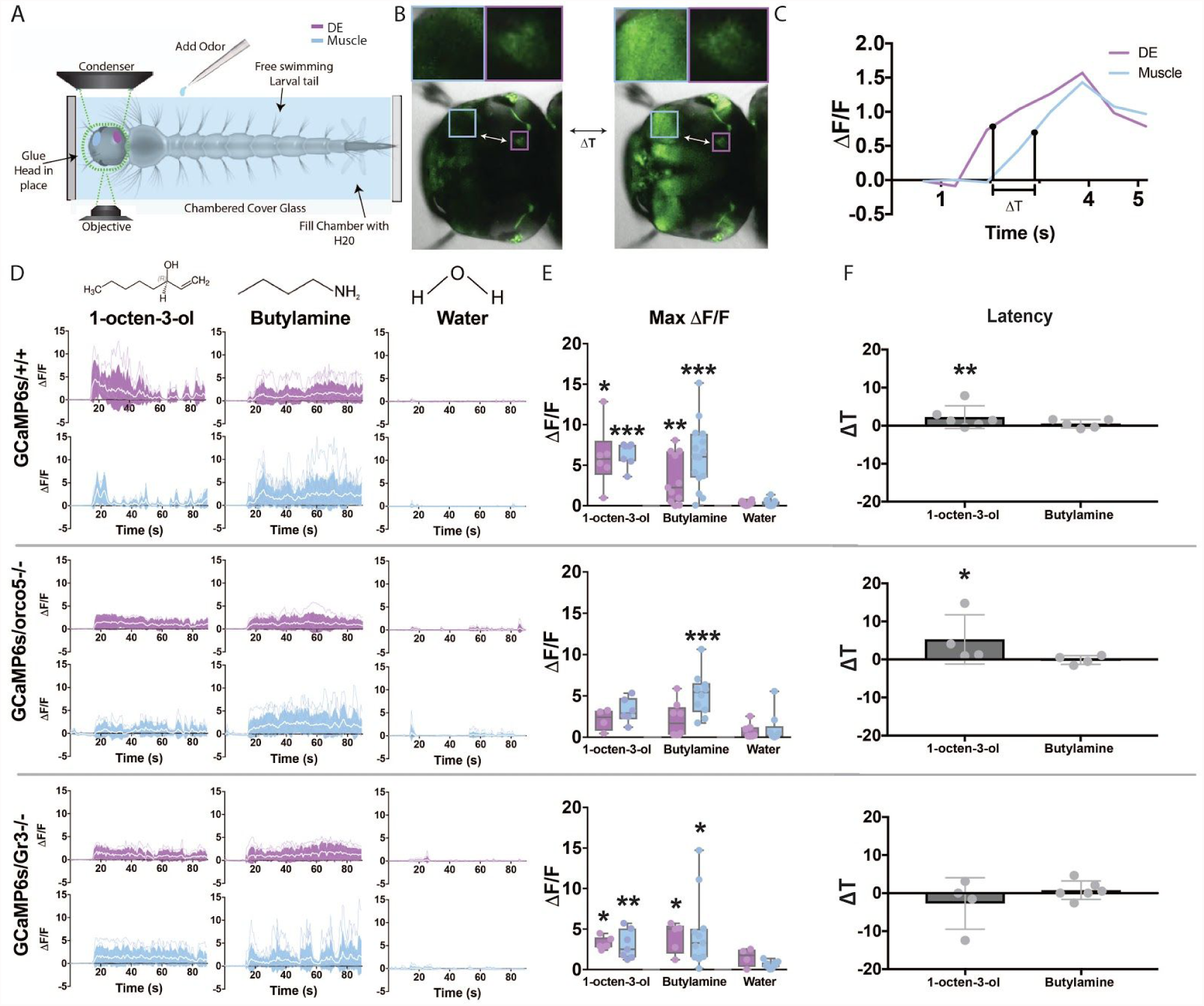
Live calcium imaging of stimulus-evoked responses in GCaMP6s/+/+, GCaMP6s/orco5-/-, and GCaMP6s/Gr3-/-. Mosquito larval calcium responses to various stimulants were recorded using a Leica SP5 Confocal microscope. To secure the larval head for imaging while allowing free movement of the larval tail, the dorsal side of the larva’s head was adhered to a chambered cover glass using quick setting adhesive. The chamber was then filled with water and the larvae was allowed to rest before being introduced to stimulants (A). To compare temporal difference between the Deuterocerebrum (DE,purple) and muscles (blue), the difference between time points at 50% of the maximum ΔF/F of the first response peak following the addition of stimuli (B,C). Stimuli, including 1-octen-3-ol, butylamine, and water were introduced to GCaMP6s/+/+, GCaMP6s/orco5-/-, and GCaMP6s/Gr3-/- larvae 15 sec after the start of recording. Calcium responses within the Deuterocerebrum and muscles were recorded at 0.645 frames/sec (D) and maximum fluorescence values were plotted (E). The temporal difference in seconds between the DE and Muscle responses were calculated and plotted by comparing DE and muscle timepoints at 50% of maximum ΔF/F (F). The number of biological replicates used for each experiment were 3 or greater. Differences in ΔF/F and Latency were analyzed using a Welch’s T-test and a Mann-Whitney U test respectively. *: p-value < 0.05, **: p-value < 0.01, ***: p-value < 0.001.

**Figure 3.**
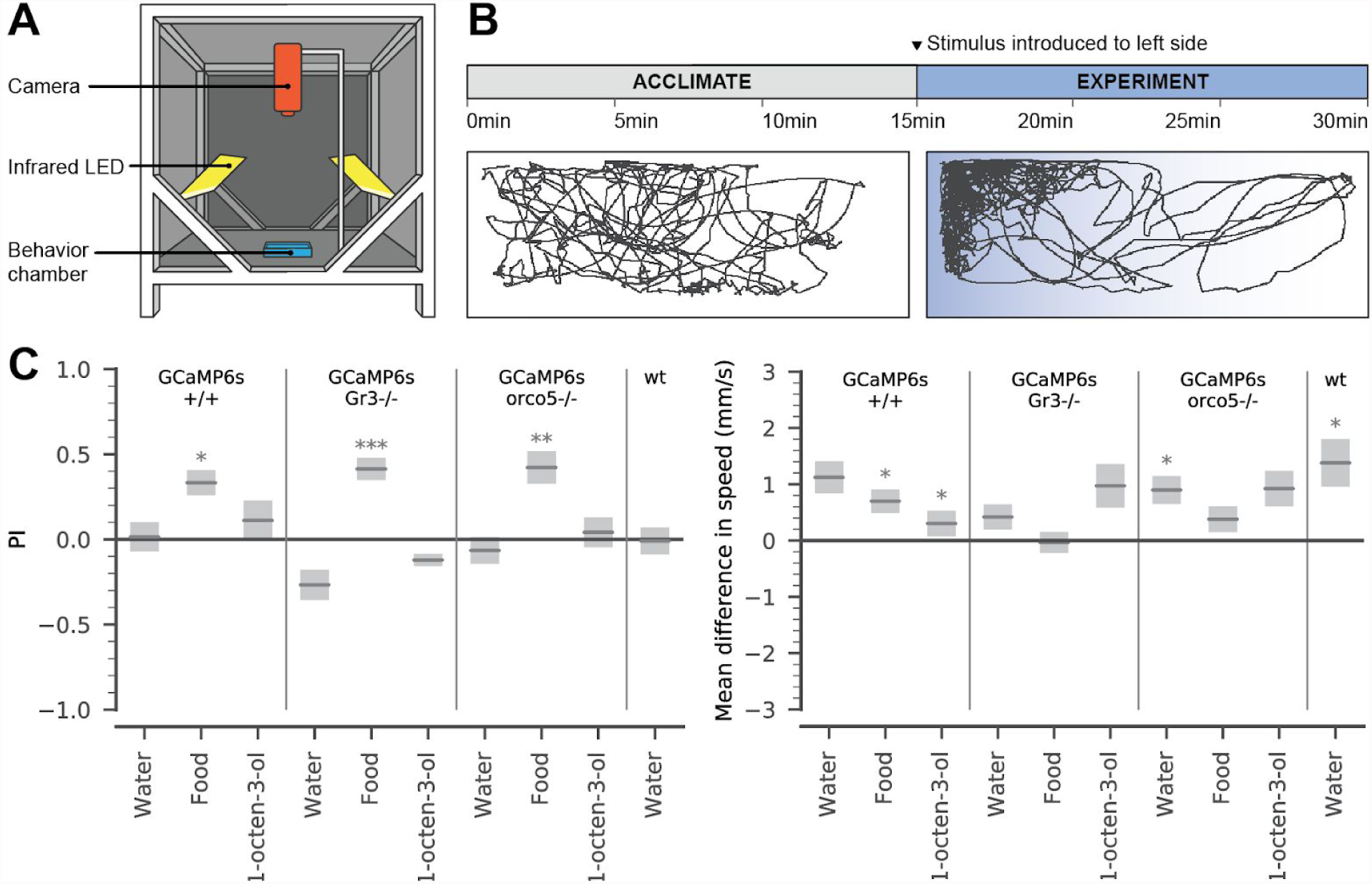
Behavioral analysis of stimulus-evoked responses in GCaMP6s/+/+, GCaMP6s/orco5-/-, GCaMP6s/Gr3-/-, and wt larvae. The dark experimental arena used for behavior testing. Animals were released individually into a custom 3D printed porcelain behavior chamber (blue), lit with infrared LED panels (yellow) and recorded with a Basler Scout Machine Vision Area Scan GigE camera (orange) (A). In each experiment, the larva was allowed to acclimate in the arena for 15 minutes. Next, 100uL of one stimulus was introduced to the upper left side of the arena. In both stages, larval behavior was recorded at 2fps, and larval position in each frame was extracted using ImageJ and Python. This example trajectory shows the movement of a GCaMP6s/orco5-/- larva before and after the addition of 100uL food extract (B). Using these trajectories, we compared PI (defined as the proportion of time spent in the odor half of the arena minus the proportion of time spent in the non-odor half) and mean speed across all larvae during the experiment phase (C). mean +/- SEM, p-values: Welch’s T-test comparing acclimation period to experiment period. *: p-value < 0.05, **: p-value < 0.01, ***: p-value < 0.001, n = 14 ~ 20 per treatment.

### Larval Brain responses to Olfactory Stimuli in Mutant Genetic Backgrounds

To gain further insight into the genetic basis for neuronal responses to various stimuli, we genetically crossed GCaMP6s/+/+ mosquitoes into two separate genetic backgrounds that harbored homozygous mutations in either an important odorant coreceptor (orco; [19]) or a subunit of the heteromeric CO_2_ receptor (Gr3-/-; [20]) (Supplementary Figure 4). Using our larval tethered-swimming imaging assay and stimulus panel described above to compare calcium evoked responses between GCaMP6s/+/+, GCaMP6s/orco5-/-, GCaMP6s/Gr3-/- enabled us to parse out receptors important for eliciting responses to various stimuli. Interestingly, we found that, compared to GCaMP6s/+/+, the DE and muscles of GCaMP6s/orco5-/- elicited fewer significant responses to stimuli (p-value > as well as a general reduction in calcium evoked responses to all stimuli (Supplemental Figures 2, 3B). Only butylamine elicited significant responses compared to the water control in the muscles (max ΔF/F0 is 5.20±2.60, p-value = 8.55e-4) (Figure 2E, Supplementary Figure 3B). Additionally, when comparing DE and muscle responses between GCaMP6s/orco5-/- and GCaMP6s+/+, only muscle responses to 1-octen-3-ol and VUAA1 within GCaMP6s/orco5-/- showed a significant decrease (Supplemental Figure 3E). A strong decrease in DE response to 1-octen-3-ol was also seen in GCaMP6s/orco5-/-, however it was not significant (max ΔF/F is 2.14 ± 1.30, p-value = 0.0563). Examining the latency of response in GCaMP6s/orco5-/- showed that stimulation with 1-octen-3-ol elicited responses in muscles 5.27±6.50 sec after activation of the DE (p-value = 0.04412), while butylamine showed little difference in the latency between DE and muscle response (0.13±1.14 sec; p-value = 0.9492)(Figure 2F). Furthermore, although the intensity DE of response to 1-octen-3-ol was not as strong as that of GCaMP6s+/+, the response was seen to persist for a longer period of time (Supplementary Figure 7). Taken together, our results demonstrates orco’s role in the detection of numerous chemosensory stimuli. Additionally, we found that orco may play an important role in 1-octen-3-ol detection and response. A nonsignificant reduction of response in the DE yet a significant reduction in the muscles indicate that even strong yet insignificant decreases in neural response may lead to a significant reduction of muscle output.

In contrast to the GCaMP6s/orco5-/- mutants, GCaMP6s/Gr3-/- mutants showed more robust calcium-evoked responses, and were generally not significantly different from those of GCaMP6s/+/+ with the exclusion of muscle responses to 1-octen-3-ol (Supplemental Figure 3C-E). For instance, relative to the water control, 1-octen-3-ol and ethyl acetate elicited strong responses in both the DE and muscle (p-value < 0.05) (Supplemental Figure 3). In total, five stimuli evoked significant increases in fluorescence within muscles of GCaMP6s/Gr3/-/- mutants; including 1-octen-3-ol (max ΔF/F0 is 3.08 ±1.79, p-value = 0.009765), butylamine (max ΔF/F0 is 4.59±4.46, p-value = 0.01383), ethyl acetate (max ΔF/F0 is 4.51±1.85, p-value = 0.001199), VUAA1 (max ΔF/F0 is 1.86 ±1.11, p-value = 0.01393), and sucrose (max ΔF/F0 is 2.03±1.47, p-value = 0.04154)(Supplementary Figure 3C). Interestingly, the latency in response between the DE and muscle ROIs were near-simultaneous for butylamine (0.78 ± 2.61 sec, p-value = 0.6565) and 1-octen-3-ol (2.72 ± 6.75 sec, p-value = 0.9389), with the latter also demonstrating more persistent responses in both the DE and muscles, suggesting that gustation or other chemosensory channels may be involved in the processing of these odorants (Figure 2F, Supplementary Figure 7).

### Odor-evoked behavior in free-swimming larvae

Previous studies have shown that mosquito larvae respond behaviorally to chemosensory stimuli including 1-octen-3-ol [24], but the genetic basis of these responses remain unclear. To investigate the behavioral responses of the GCaMP6s larvae in various genetic backgrounds (GCaMP6s/+/+, GCaMP6s/ocro5-/-, and GCaMP6s/Gr3-/-), we examined free-swimming larval responses to a limited odor panel in a custom arena. Individual larvae were allowed to acclimate inside the dark behavior arena before a stimulus - either food extract, 1-octen-3-ol, or a water-only (negative) control - was added to one side of the arena, and responses were analyzed and compared for the 15-minute acclimation period and the following 15-minute experiment period (Figure 3, n > 14 for all treatments). From the videos, we were able to quantify a variety of behavioral responses, including the larva’s preference index (PI, defined as the proportion of time spent in the odor half of the arena minus the proportion of time spent in the non-odor half), and mean swimming speed (mm/s), as well as other kinematic variables (Supplemental Figure 5). Importantly, prior to stimulation we found no differences in position or mean speed between larvae of the mutant backgrounds, and between the mutants and the Liverpool strain background (n= 1-way ANOVA p-value = 0.315 [PI]; p-value = 0.3173 [mean speed], Supplemental Figure 5). In all strains, the addition of water had no significant influence on which side of the chamber the larvae preferred (p-values > 0.12, Welch’s t-test compared to acclimation period). Results from these experiments showed that the larvae of all strains significantly preferred the side of the chamber with the food extract (p-values < 0.05, Welch’s t-test compared to acclimation period), and this preference was consistent across all strains (p-value = 1 for all strain comparisons, pairwise t-test with Holm-Bonferroni correction). Although larvae of all three strains showed no significant positional preference for 1-octen-3-ol (p-values > 0.14, Welch’s t-test compared to water control on the same strain; p-values > 0.13, t-test compared to acclimation period), GCaMP6s/+/+ larvae significantly decreased their speed compared to the water stimulus (p-value = 0.041, pairwise *t*-test).

### Discussion

In these experiments, we have expanded the toolbox of techniques for investigating a globally important disease vector, *Aedes aegypti*, and demonstrated the versatility of these tools for investigating overarching questions in neurobiology such as sensory integration and information processing. Furthermore, our results provide novel insights into *Ae. aegypti* chemosensory processing that may help guide future research in disease-vector mosquito control.

The robust expression of GCaMP6s in various mosquito tissues (Figure 1) allows quantification of stimulus-evoked responses in both motor and sensory systems, and in the adult and larval stages, including the adult antennae, adult maxillary palps, larval deutocerebrum (DE), and larval muscle (Figure 2, Supplemental Figure 1). This broad GCaMP6s expression allowed us to investigate both motor and sensory responses in *Ae. aegypti* larvae to an ecologically relevant panel of chemosensory stimuli. These cues elicited spatiotemporal patterns in GCaMP6/+/+ larval muscle and central nervous system (CNS), and revealed that key components of natural odors may be relevant across *Ae. aegypti* life stages (Supplemental Figure 2). Further, when we crossed this GCaMP6s/+/+ line with orco5-/- mutant to generate GCaMP6s/orco/-/-, we observed attenuation in these stimulus-evoked responses, particularly in response to known OR ligands (VUAA1 [42], 1-octen-3-ol [43], Supplemental Figure 3). By contrast, GCaMP6s/Gr3-/- larvae showed no significant impairment in response to any of the odorants tested, indicating that the heteromeric CO_2_ receptor complex is not critical for the detection and response to these stimuli in the larval stage. This supports previous transcriptome work suggesting that the Gr3 receptor is expressed at very low levels in *Ae. aegypti* larvae [44]. Together, these results demonstrate the utility of these GCaMP6 mutants for investigating the neural representation of chemosensory-mediated stimuli. Interestingly, our neuronal imaging showed no significant response to food odors, however muscle responses were observed. This may reflect the fact that our dorsal imaging plane did not extend into the ventral sub-oesophageal ganglion (SOG), which is innervated by sensory nerves from the mouthparts. Finally, our behavioral experiments contextualized some of these stimulus-evoked responses in a more naturalistic environment, revealing that ORs may act in parallel with other chemosensory channels during foraging behavior in *Ae. aegypti* larvae (Figure 3). Together, our combination of calcium imaging and behavior experiments highlights the importance of studying chemosensory behavior from multiple perspectives, and build on earlier work on the chemosensory repertoire [45,46] and behaviors of mosquito larvae [47,48] to gain a more complete understanding of mosquito chemical ecology.

Our results highlight important avenues of future research in mosquito sensory processing. First, the mechanisms of chemosensory cue detection in *Ae. aegypti* larvae remains an open question. In terrestrial environments, long-range chemosensory stimuli are largely limited to volatile compounds with high vapor pressure at ambient temperatures [49]. However, *Ae. aegypti* larvae inhabit an aquatic environment that is a rich source of chemical signals far more varied in size, polarity, and structure, such as large proteins, amino acids, long hydrocarbon chains, and multi-molecular fragments of organic debris [50]. Interestingly, *Ae. aegypti* larvae express far fewer ORs than adults [45] and have a markedly smaller and physiologically less developed antennal lobe [51,52](Supplemental Figure 5). *Ae.aegypti* larvae may rely on a more diverse assortment of IRs and GRs, in addition to ORs, to detect the wide range of water-borne chemicals relevant to behaviors such as foraging and predator avoidance [26], [53]).

Characterization of the *Ae. aegypti* larval IRs and GRs may help identify chemical compounds that are most relevant to larval environments, and lend insight into the spectrum of larval chemical receptors. In addition to receptor-level chemical detection, the mechanism of chemosensory processing in the *Ae. aegypti* larval CNS is not well understood. Aquatic crustaceans integrate information from hydrodynamic detectors and two distinct types of chemosensory receptors within the CNS [54,55], but it is unclear if *Ae. aegypti* sensory transduction follows this same model. From an evolutionary perspective, comparing the mechanism of *Ae. aegypti* larval olfaction to crustacean, amphibian, and fish models may also provide critical insight into the convergent evolution of aquatic chemosensation. Our *Ae. aegypti* GCaMP6/+/+ strain is of particular interest as the first example of GCaMP6 expression in an aquatic insect model.

Additionally, some odor components are shared among multiple ecologically relevant cues for mosquitoes, and the neurobiological implications of these correlations are unclear. For example, 1-octen-3-ol is a component of both host odors [56] and microbial byproducts [57] that may function as food for larval mosquitoes. It is not unreasonable to hypothesize that there may be strong evolutionary selection on mosquito ORs that are beneficial in both life-history stages, and if so, identifying those chemicals that operate as both larval attractants and adult host cues may provide attractants that can be leveraged for mosquito control. Moreover, the mechanism of chemotaxis in *Ae. aegypti* larvae remains an open question. In other insect models such as *D. melanogaster*, larvae employ active sampling strategies to locate and navigate to food cues [58]. But it is unclear how *Ae. aegypti* larvae navigate chemosensory signals in an aquatic environment that is quite different in volume and turbidity from those experienced by *D. melanogaster* or even *E coli [59]* and *C. elegans [60]*, which navigate chemosensory gradients at a significantly smaller scale. Quantitative modeling and further behavioral experiments may help better understand chemotaxis in an enigmatic aquatic insect model, and highlight interesting commonalities and differences in navigation strategy across different environments and spatial scales.

Generalizing further, our GCaMP6s/Gr3-/- and GCaMP6s/orco5-/- mutants could address gaps in our broader understanding of multisensory integration and sensorimotor responses, particularly in adult mosquitoes. Behavioral work in *Ae. aegypti* adults presents compelling evidence for the involvement of multisensory integration in host-seeking [61,62]. However, little is known about the neural bases of these behaviors. In *D. melanogaster*, GCaMP6s imaging has revealed the functional basis of information convergence in higher-order brain areas [63,64]. Future work with our GCaMP6s/+/+ line may similarly help decode the neural representations of multimodal host cues in mosquitoes, and provides motivation for the development of transcriptional control systems such as GA4/UAS or the Q-system [65] for tissue-specific GCaMP6s expression. Importantly, we observed high GCaMP6s expression in both muscle and neuropil (Figure 2, Supplementary Figure 6). In *D. melanogaster*, concurrent analysis of neural response and motor output has facilitated experiments in the integration of sensory processing and sensory-motor transformations [66–68]. By taking advantage of this simultaneous recording capacity in *Ae. aegypti* mosquitoes, additional experiments could investigate how these multisensory integration pathways mediate motor responses, and ultimately, determine behavioral decisions such as host choice and oviposition site preference. Finally, these GCaMP6s/+/+ mosquitoes and GCaMP6s/Gr3-/- and GCaMP6s/orco5-/- mutants could provide additional information and strategies for the control of disease-vector mosquitoes. Female mosquitoes may use olfactory indicators of larval habitat quality to choose oviposition sites [69]. A better understanding of chemosensory cues that elicit strong responses in larvae could help identify new attractants for use in oviposition traps, or oviposition deterrents for use in homes and outdoor water containers.

## Materials and Methods

### Insect rearing

Mosquitoes used in all experiments were derived from of the *Ae. aegypti* Liverpool strain, which was the source strain for the reference genome sequence. Mosquitoes were raised in incubators at 28°C with 70–80% relative humidity and a 12 hour light/dark cycle. Larvae were fed ground fish food (TetraMin Tropical Flakes, Tetra Werke, Melle, Germany) and adults were fed with 0.3M aqueous sucrose. Adult females were blood fed three to five days after eclosion using anesthetized mice. All animals were handled in accordance with the guide for the care and use of laboratory animals as recommended by the National Institutes of Health and supervised by the local Institutional Animal Care and Use Committee (IACUC).

### Construct Assembly

To generate the GCaMP6s plasmid (plasmid sequence and DNA available for order at addgene ID# 106868), components were cloned into the *piggyBac* plasmid pBac-3xP3-dsRed [70] using Gibson assembly/EA cloning [71]. Specifically, pBac-3xP3-dsRed was digested with BstBI and SacII, and an attB site, amplified from a stock attB plasmid [72] with primers 997.C5 and 997.C6. The predicted *Ae. aegypti polyubiquitin* (*PUb*) promoter fragment [73] was amplified from *Ae. aegypti* genomic DNA using primers 997.C1 and 997.C2. While the GCaMP6s fragment [30] was amplified from vector pGP-CMV-GCaMP6s (Addgene plasmid #40753) using primers 997.C3 and 997.C4 and cloned in via EA cloning. The resulting plasmid was then digested with PacI and AvrII and the following fragments cloned in via EA cloning. P10 3’UTR [74] was amplified with primers 997.C7 and 997.C8 from vector pJFRC81-10XUAS-IVS-Syn21-GFP-p10 (Addgene plasmid 36432) and the OpIE-2 promoter region [75] amplified from vector pIZ/V5-His/CAT (Invitrogen) using primers 997.C9 and 997.C10. The plasmid was grown in strain JM109 chemically competent cells (Zymo Research #T3005) and isolated using Zyppy Plasmid Miniprep (Zymo Research #D4037) and maxiprep (Zymo Research #D4028) kits. The full plasmid sequence was verified using Source Bioscience Sanger sequencing services. A list of primer sequences used in the above construct assembly can be found in Supplementary Table 1.

### Generation of GCaMP6s/+/+, GCaMP6s/Gr3-/-, and GCaMP6s/orco5-/- transgenic lines

GCaMP6s/+/+ mosquitoes were created by injecting 0-1hr pre-blastoderm stage embryos with a mixture of the GCaMP6s plasmid described above (200ng/ul) and a source of piggyBac transposase (phsp-Pbac, (200ng/ul))[76–79]. Embryonic collection and microinjections were largely performed following previously established procedures [70,80]. Injected embryos were hatched in deoxygenated water and surviving adults were placed into cages. Adult G0 females were allowed a blood-meal 4 days after eclosion. Following general rearing procedures described above, G1 larvae were screened for expected fluorescent markers, OpIE-2-dsRed, and *PUb-*GCaMP6s (Figure 1C, Supplementary Figure 4D). Larvae with positive fluorescent signals were collected under a fluorescent stereomicroscope (Leica M165FC). To ensure that this line was produced with a single chromosomal insertion, single individuals from each of the lines were backcrossed for four generations to our wild-type stock. Mendelian transmission ratios for each generation were measured. In all cases, we observed a 50% transmission ratio in each generation, indicating that our strain represented an insertion on a single chromosome. To obtain a nearly complete homozygous line, our GCaMP6s line was screened and selected for at least 20 generations. To obtain the GCaMP6s/orco5-/- homozygous line, GCaMP6s/+/+ (♂) was crossed with orco5-/- (♀), then G1 individuals (♂) with the GCaMP6s phenotype were backcrossed with orco5-/- individuals (♀) for at least 8 generations as single mosquito pairwise matings, sanger sequencing was utilized to confirm GCaMP6s/orco5-/- homozygosity (Supplemental Figure 4A,B). To obtain the GCaMP6s/Gr3-/- mutant homozygous line, GCaMP6s/+/+ (♂) was crossed with GR3-/- (♀, labeled with a CFP marker), then continually selected for individuals with correct markers (dsRed, GCaMP6s, and CFP). Furthermore, single mosquito pairwise crosses were performed for at least 8 generations (Supplementary Figure 4C,D). The transgenic GCaMP6s/+/+ line has been deposited at BEI MR4 Resources (Accession # still waiting for acceptance of strain from BEI MR4).

### Genetics and Molecular Characterization of Insertion Site

To characterize the insertion site of GCaMP6s, we modified an inverse PCR protocol described previously [70,81]. Briefly, genomic DNA(gDNA) was extracted from 10 *Ae. aegypti* fourth instar larvae using the DNeasy Blood & Tissue Kit (Qiagen, Hilden, Germany) in accordance with the manufacturer’s protocol. The eluted DNA was diluted, and two separate restriction digests were performed to characterize both the 5’ and 3’ ends using Sau3AI (5’ reaction) or HinP1I (3’ reaction) restriction enzymes. A ligation step using NEB T4 DNA Ligase was then performed on the restriction digest products to promote circularization of digested DNA. Two rounds of PCR were performed using primers 991.5F1, 991.5R1, 991.5F2, 991.5R2, 991.3F1, 991.3R1, 991.3F2 and 991.3R2 (with their corresponding restriction digest reaction) and sequence confirmation (1018) are listed in Supplementary Table 2. PCR products from the second round of PCR were cleaned using the MinElute PCR Purification Kit (Qiagen) in accordance with the manufacturer’s protocol, and subsequently sequenced by Sanger sequencing (Source BioScience). Both the location and orientation (chromosome 2, with the flanking genomic regions for the 5’ and 3’ *piggyBac* ends positioned at the genomic loci 285,175,805-285,176,289 and 285,175,275-285,175,803, respectively) were confirmed by PCR using primers designed from the mapped genomic region in combination with both 3’ piggyBac end forward primers. Sequencing data was then blasted to the AaegL5.0 reference genome. Alignment of the sequencing data was performed using EMBOSSWater (https://www.edi.ac.uk/Tools/psa/emboss_water/).

### Odor-Evoked Confocal Imaging of Non-water submerged Larvae/Adult

For larval imaging of GCaMP6s/+/+ calcium transients, using a slightly moistened fine tip paint brush, larva were placed ventral side down on double-sided tape adhered to a clean glass slide. Due to the larvae being exposed to air rather than its normal aquatic environment, to prevent dessication a moistened fine tip paint brush was used to periodically wet the larvae without affecting the sticky-tape adhesive. Imaging was focused on the full body and head. For adult imaging, mosquitoes were placed laterally on double-sided tape after being placed on ice for approximately 10 minutes. Antennae and proboscis were immobilized by using an artist brush and gently brushing the respective appendages onto the double sided tape. For both larvae and adults, a minimum of 15 seconds of inactivity was first captured recording the specimen. Recording continued for an additional 35 seconds. Images and recordings were taken using an Inverted Confocal microscope (Leica SP5).

### Odor-Evoked Confocal Imaging of Larvae in Tethered-Swimming Assay

To immobilize each larval head, while allowing for movement of the larval body, less than one microliter of clear Aron Alpha high strength rapid bonding adhesive (Catalog # 72588) was applied to a Lab-Tek II chambered #1.5 German coverglass system composed of transparent borosilicate glass (Thermo Catalog #155382). Immediately following the application of the adhesive, the ventral side of a single larva was placed directly onto the adhesive, rapidly bonding the larval head to the coverglass in less than 1 minute. The chamber was then filled with 500 uL of deionized water to fully submerge the larvae, while allowing for the larva’s respiratory siphon to meet the surface of the water. Before any recordings, the larvae was allowed to rest for 12 hours to assimilate to the preparation. Recordings of stimulus-induced fluorescent responses were taken around the head. 100 µL of 5% solution of odorants were injected into the chamber after 15 seconds of inactivity in larval brains. Activity was measured from 15 seconds prior to addition of stimulus to 90 seconds after. Following each trial, stimuli were removed by draining the water in chamber, gently flushing the larvae and chamber three times, and refilling with fresh deionized water. The same larva was used for multiple stimulants.

### Selection and preparation of Odorants

Stimulants were chosen from a list of known olfactory and/or gustatory stimuli of both *Drosophila melanogaster* and adult *Ae. aegypti [24,25,33–38]*. These included ethyl acetate (Sigma Cat# 319902), lactic acid (Sigma Cat# L1750), 1-octen-3-ol (Sigma Cat# O5284), butylamine (Sigma Cat# 471305), VUAA1 (Vitas-M Cat# STK047588), sucrose (Sigma Cat# S0389), lobeline (Sigma Cat# 141879), glutamate (Sigma Cat# 49621), and water (negative control). All stock solutions of odors were prepared as 5% solutions in water, with a final bioassayed concentration of 6×10^−5^M. Food extract for larval experiments was prepared by mixing 0.5% fish food (Hikari Tropic First Bites: Petco, San Diego, CA, USA) in milliQ water. The solution was allowed to sit for one hour, then filtered through a 0.2 μm sterile filter (#28145-477, VWR International, Radnor, PA, USA) to remove solid particulates. 1-octen-3-ol used in behavior experiments was prepared as a 10^−4^M solution in water.

### Confocal Imaging/Data analyses

To quantify fluorescence responses to various stimuli, Leica LAS X Core Offline version 3.3.0 software was used to export raw fluorescence data from relevant ROIs. Further analysis was done using GraphPad Prism and RStudio. To account for differences in fluorescence intensity that differed between each larva, raw fluorescence was normalized using ΔF/F_0_= (F-F_0_)/F_0_ where F is mean intensity of fluorescence at a certain time point and F_0_ is the baseline level of fluorescence using the average fluorescence intensity from the first 15 seconds of the recording without stimulation [82]. To compare the differences between GCaMP6s/+/+, GCaMP6s/orco5-/-, and GCaMP6s/Gr3-/- calcium responses to our simulus panel, a Welch’s t-test was conducted comparing the ΔF/F_0_ of multiple replicates between the two larval backgrounds in response to the same stimulus. Importantly, due to the methodology of our larval imaging assay, the larval abdomen would occasionally be viewable behind the ROI. To confirm that this interference does not create any significant artifacts while measuring raw fluorescence, raw fluorescence 2 seconds before and during interference were compared and no significant interference was detected (t = 0.237, p-value = 0.8158).

### Muscle/DE Latency Analysis

Temporal differences between muscle and DE responses were calculated by subtracting DE timepoints at 50% of maximum ΔF/F_0_ of the first peak following the addition of stimulus from that of muscles (Supplementary Figure 2B,C). Recordings with both DE and muscles not displaying clear peaks in response to stimulants as well as latency values greater than 15 were treated as NA. Latency values were converted into ordinal values of 4 categories: NA, negative, 0 (no difference), and positive. A Mann-Whitney test between latency values from each stimulus and that of water was used to determine significant differences.

### Free-Swimming Larval Behavior Experiments

Larvae used for free-swimming behavior experiments were reared on Hikari Tropic First Bites (Petco, San Diego, CA, USA) under a 12 hour light/dark cycle. One day before the experiment, 5-day old larvae were isolated into individual Falcon™ 50mL conical centrifuge tubes (Thermo Fisher Scientific, Waltham, MA, USA) containing ~15mL milliQ water and no food. During the experiment, individual larvae were introduced to the center of a dark behavior arena developed for assaying mosquito larval chemosensory preference (Figure 3). No light was detected inside the arena under experimental conditions (LI-250A Lightmeter, instantaneous measurements, sensitive up to 0.01 umol s-1 m-2 per uA. LI-COR Biosciences #Q40129). Animals were allowed to acclimate for 15 minutes in a custom 3D printed porcelain behavior chamber (ID #XWEEPACQA, Shapeways, New York, NY, USA) containing 20mL of milliQ water. 100uL of a chemical stimulus was then pipetted into the left side of the arena. Larvae were tested only during the day phase of the diurnal light cycle.

Larval movement was recorded at 2fps for the 15 minute acclimation period, as well as the 15 minute experiment following stimulus introduction, using a Basler Scout Machine Vision Area Scan GigE camera (scA 1000-30gm, Ahrensburg, Germany) and Basler pylon Viewer Windows software. Larval trajectories were analyzed using ImageJ Fiji[83] and custom software written in Python (http://www.python.org): Multitracker by Floris van Breugel (https://github.com/florisvb/multi_tracker), as well as a batch-processing Multitracker add-on, Multivideo Multitracker by Eleanor Lutz (https://github.com/eleanorlutz/multivideo_multitracker) (Figure 3). In brief, videos were cropped and contrast-enhanced in ImageJ Fiji. Larval position was extracted using frame-by-frame subtraction in Multitracker. Trajectories were manually inspected in the Multitracker GUI, where missing data points were added and extraneous tracked objects removed. We then converted trajectory position from pixel values to mm using the ratio of the known width of the behavior container. Finally, we calculated the instantaneous speed of the larva, in mm, for each frame. Using these position and speed values, we then calculated the mean instantaneous speed (mm/s) and preference index (PI; proportion of time spent in the odor half - proportion of time spent in the non-odor half).

## Acknowledgements

This work was supported in part by NIH grants 5K22AI113060 and 1R21AI123937 to O.S.A, and 1RO1DCO13693-04 to J.A.R; National Science Foundation grants IOS-1354159 to J.A.R and DGE-1256082 to E.K.L; Air Force Office of Sponsored Research under grant FA9550-16-1-0167 to J.A.R; and the University of Washington Robin Mariko Harris Award to E.K.L. We thank Christian Bowman for help with inverse PCR, Tjinder Grewal for assistance with larval behavior assays, and Gabriella Wolff for her contribution of confocal images for Supplemental Figure 6.

## Footnotes

^*^M.B, J.S, and E.K.L contributed equally to this work. ^†^To whom all correspondence should be addressed

## Author Contributions

O.S.A and J.A.R conceived and designed experiments. J.S., E.K.L., T..Y, M.L., M.B., K.T., R.A., and A.B. performed molecular, behavioral and genetic experiments. All authors contributed to the writing, analyzed the data, and approved the final manuscript.

## Disclosure

The authors declare no competing financial interests.

**Supplementary Figure 1.**
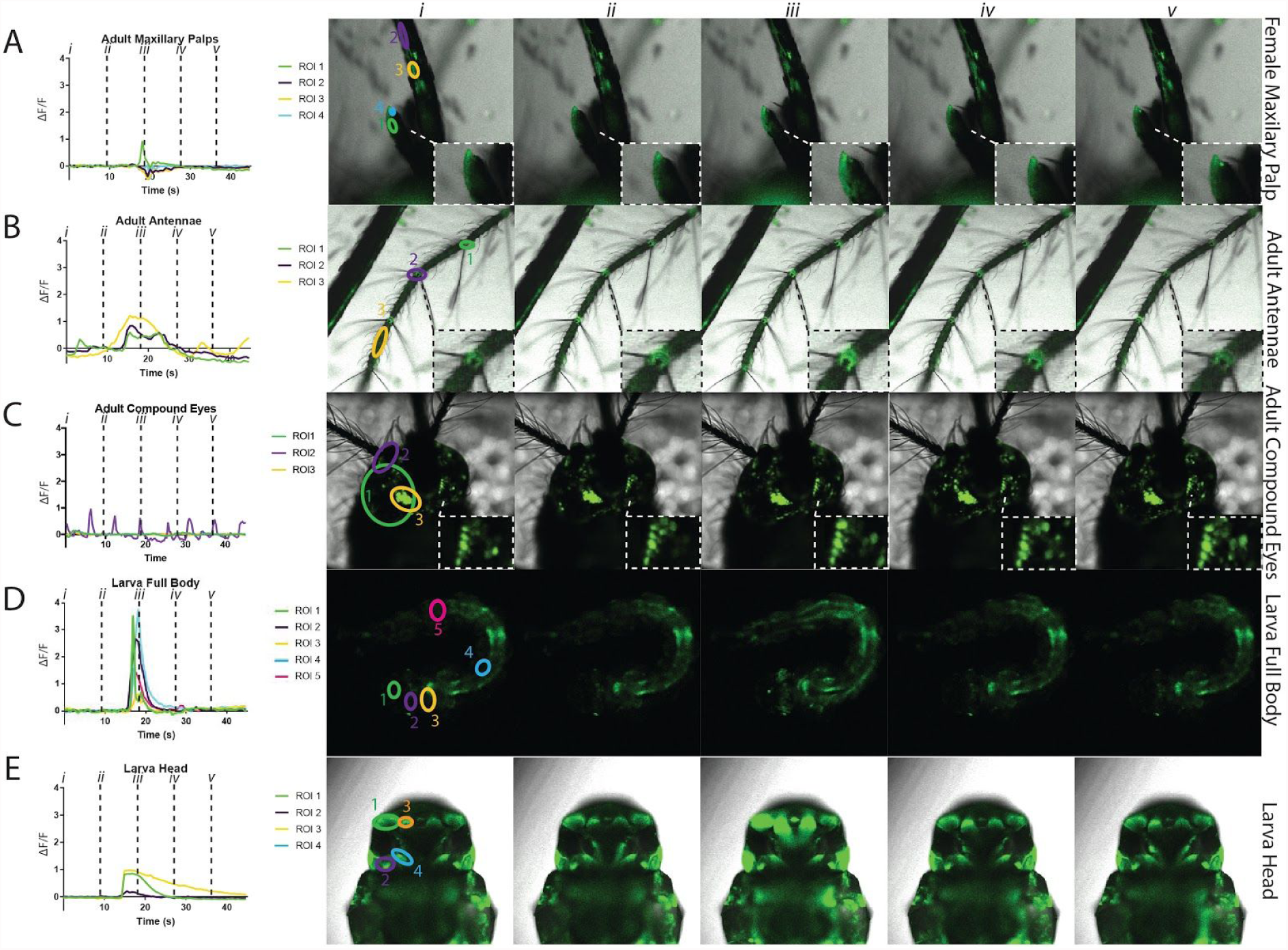
GCaMP6s is a general tool to record calcium responses in multiple tissues. Calcium responses were imaged in multiple tissues (A-E, *left*) at varying developmental stages. Frames were taken every 9s, starting at 0s (i) until 36s (v) (A-E, *right*). Calcium transients were seen in regions within the adult female maxillary palp (ROI 1,4) and labium (ROI 2,3) (A). The female adult antennae also exhibited calcium transients within the nodes (ROI 1,2) and internodes (ROI 3) of the antennal flagellum (B). Stochastic patterns of fluorescence were seen when looking at clusters of ommatidia (ROI 1-3) within the right adult compound eye (C). Calcium transients were also visualized throughout the 2nd instar larvae. For example, responses were recorded in the medial retractor muscles (ROI 1), brain (ROI 2), longitudinal muscles in the thorax (ROI 3), longitudinal muscles in the 3rd abdominal segment (ROI 4), and muscles and neurons in the 6th abdominal segment (ROI 5) (D). When imaging the dorsal side of the larval head, calcium transients were visible in the transverse retractor (ROI 1), optic lobe (ROI 2), medial retractor (ROI 3), and antennal lobe (ROI 4) (E). Full videos have been provided in the supplement (Supplemental videos 1-5).

**Supplementary Figure 2.**
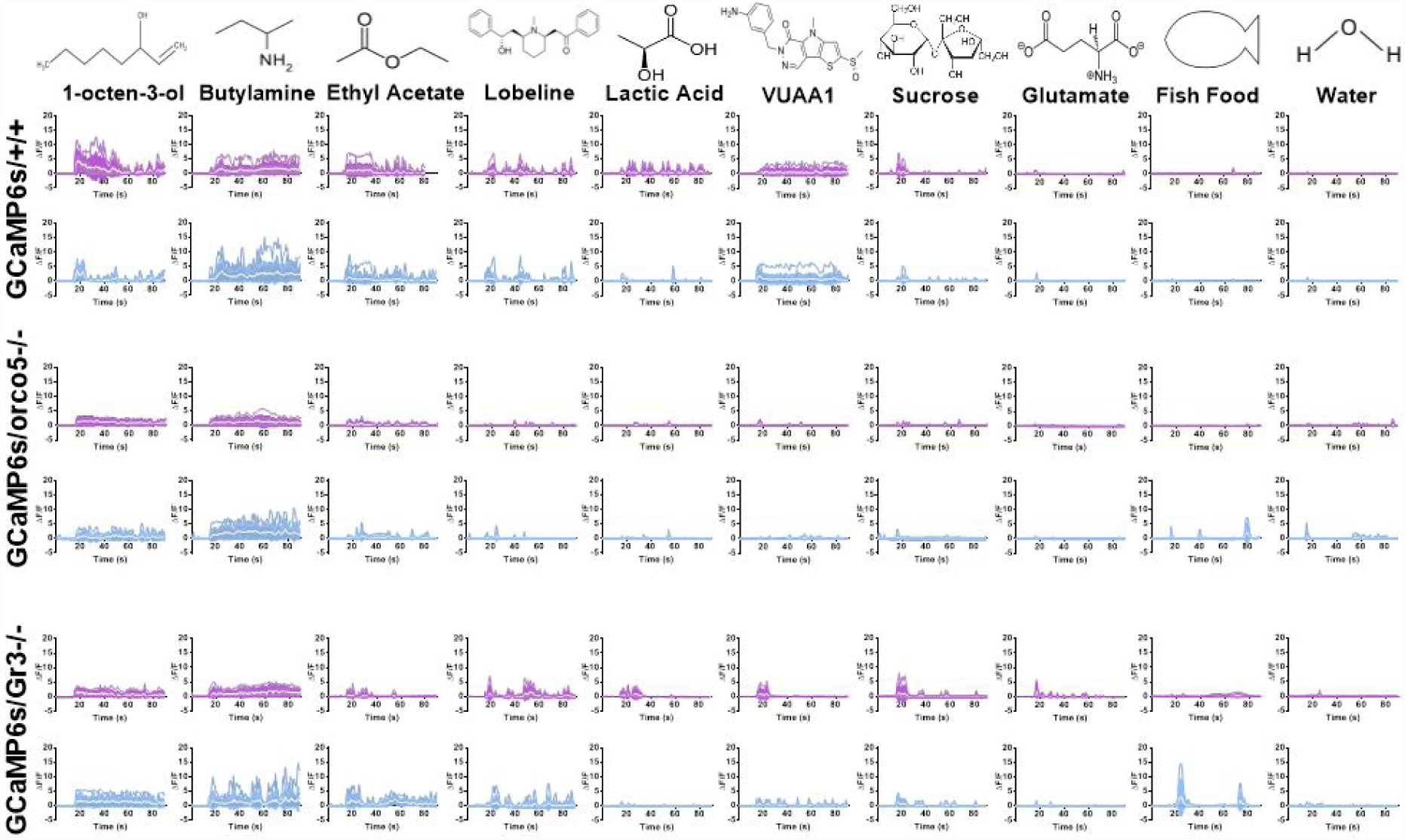
Calcium responses of GCaMP6s/+/+, GCaMP6s/orco5-/-, GCaMP6s/Gr3-/- to various stimulants. Time courses for GCaMP6s+/+, GCaMP6s/orco5-/-, and GCaMP6s/Gr3-/- DE (purple) and muscle (blue) responses to a stimulus panel including 1-octen-3-ol, butylamine, ethyl acetate, lobeline, lactic acid, VUAA1, sucrose, glutamate, fish food, and water (control). The number of biological replicates used for each experiment were 3 or greater.

**Supplementary Figure 3.**
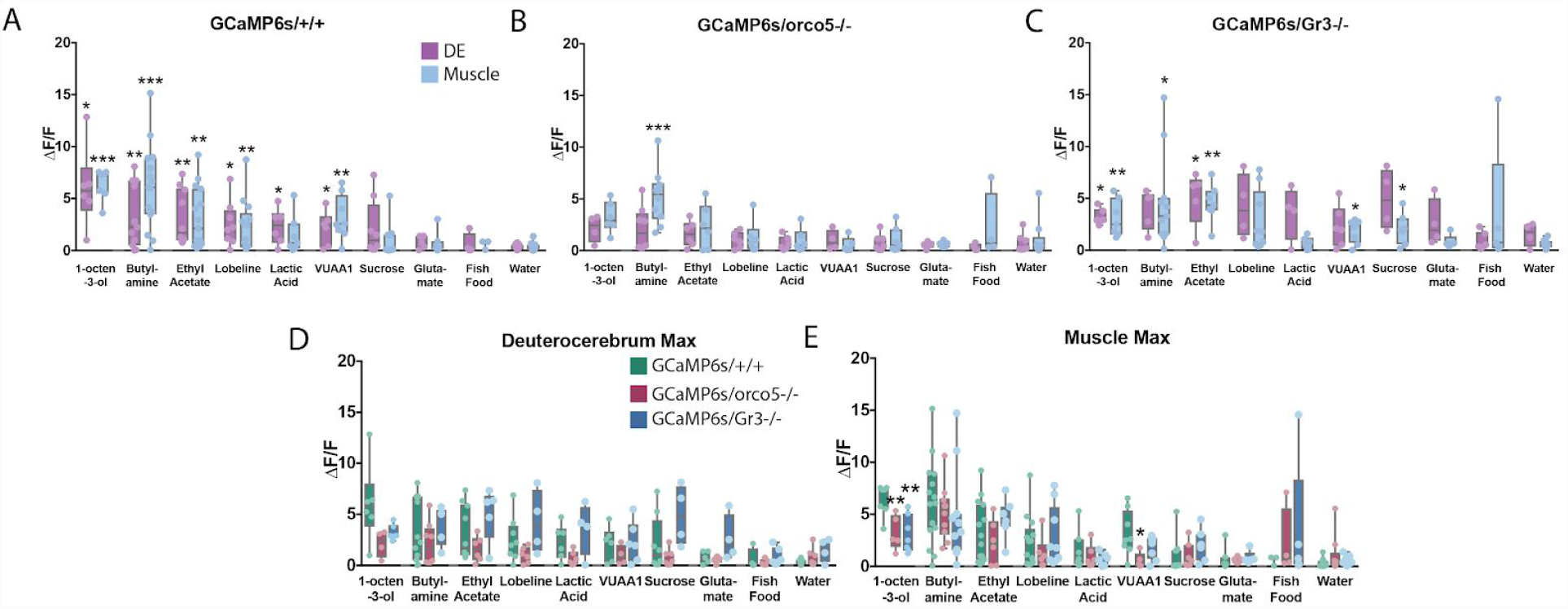
Analysis of stimuli-evoked responses of GCaMP6s/+/+, GCaMP6s/orco5-/-, GCaMP6s/Gr3-/-. Maximum fluorescence values of the DE (purple) and Muscle (blue) in response to each stimulus was compared to that of water (control) to test for significance (A-C). Maximum changes in fluorescence in response to each stimulus was also compared between the DE and muscles of GCaMP6s/+/+ (green) and both GCaMP6s/orco5-/- (red) and GCaMP6s/Gr3-/- (blue) (D,E). The number of biological replicates used for each experiment were 3 or greater. *: p-value < 0.05, **: p-value < 0.01, ***: p-value < 0.001, Welch’s T-test.

**Supplementary Figure 4.**
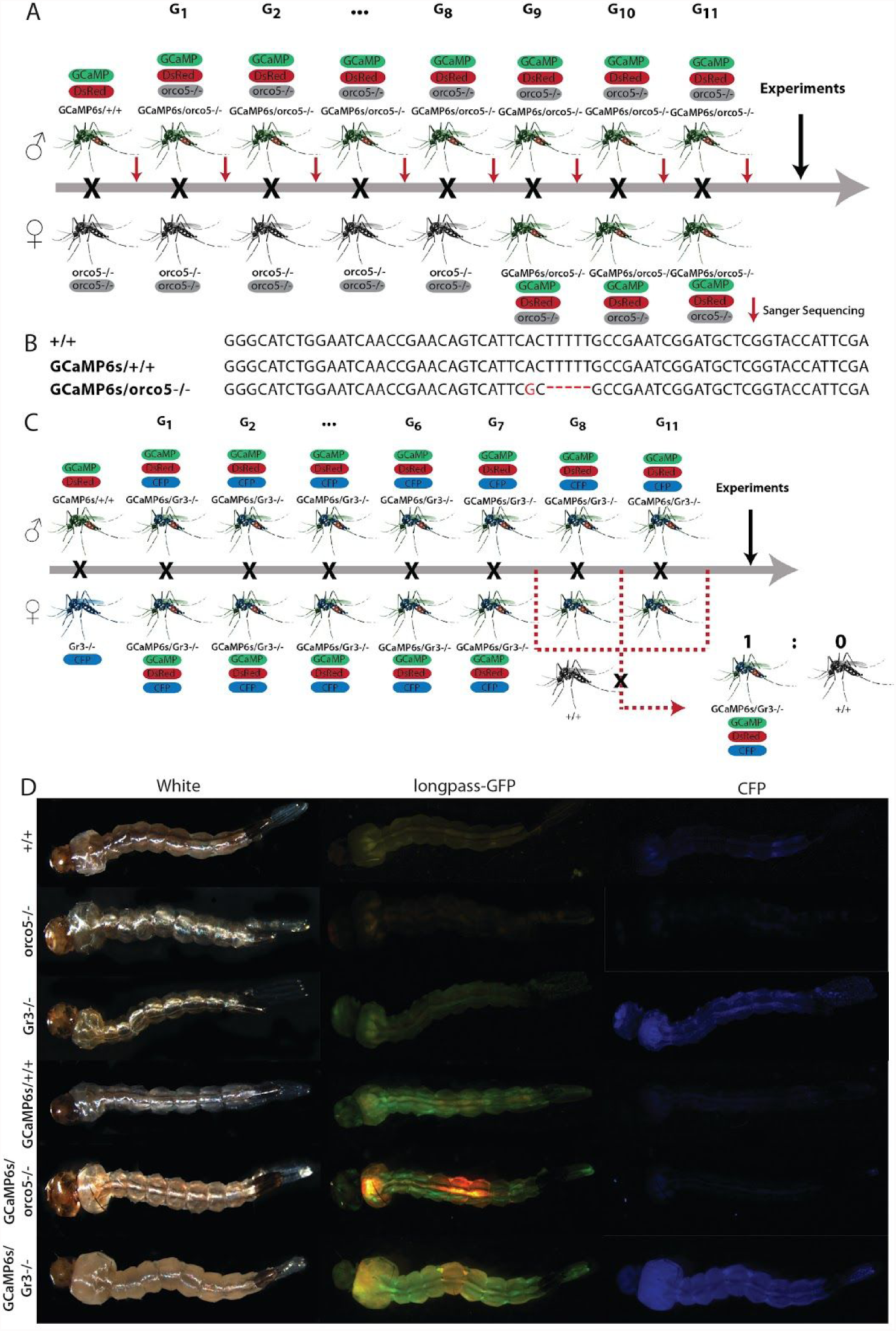
GCaMP6s/orco5-/- and GCaMP6s/Gr3-/- Line Generation and Confirmation. GCaMP6s+/+ mosquitos were mated to orco5-/- mosquitoes. Resulting individuals were mated to orco5-/- mosquitoes in a single pairwise cross for at least 8 generations and sequenced using Sanger Sequencing to confirm the presence of the *orco* gene (A). Mutations are indicated in red (B). GCaMP6s/+/+ mosquitos were crossed with Gr3-/- mosquitos. Individuals containing both GCaMP6s and Gr3-/- markers, dsRed/GFP transients and CFP respectively, were crossed in single pairwise matings for at least 8 generations to generate homozygous lines, individuals were then crossed to +/+ to confirm homozygosity by mendelian inheritance (C). All mosquito lines used were screened and sorted during the larval stage using a longpass-GFP and CFP filter to confirm OpIE-DsRed/GCaMP and CFP respectively (D).

**Supplementary Figure 5:**
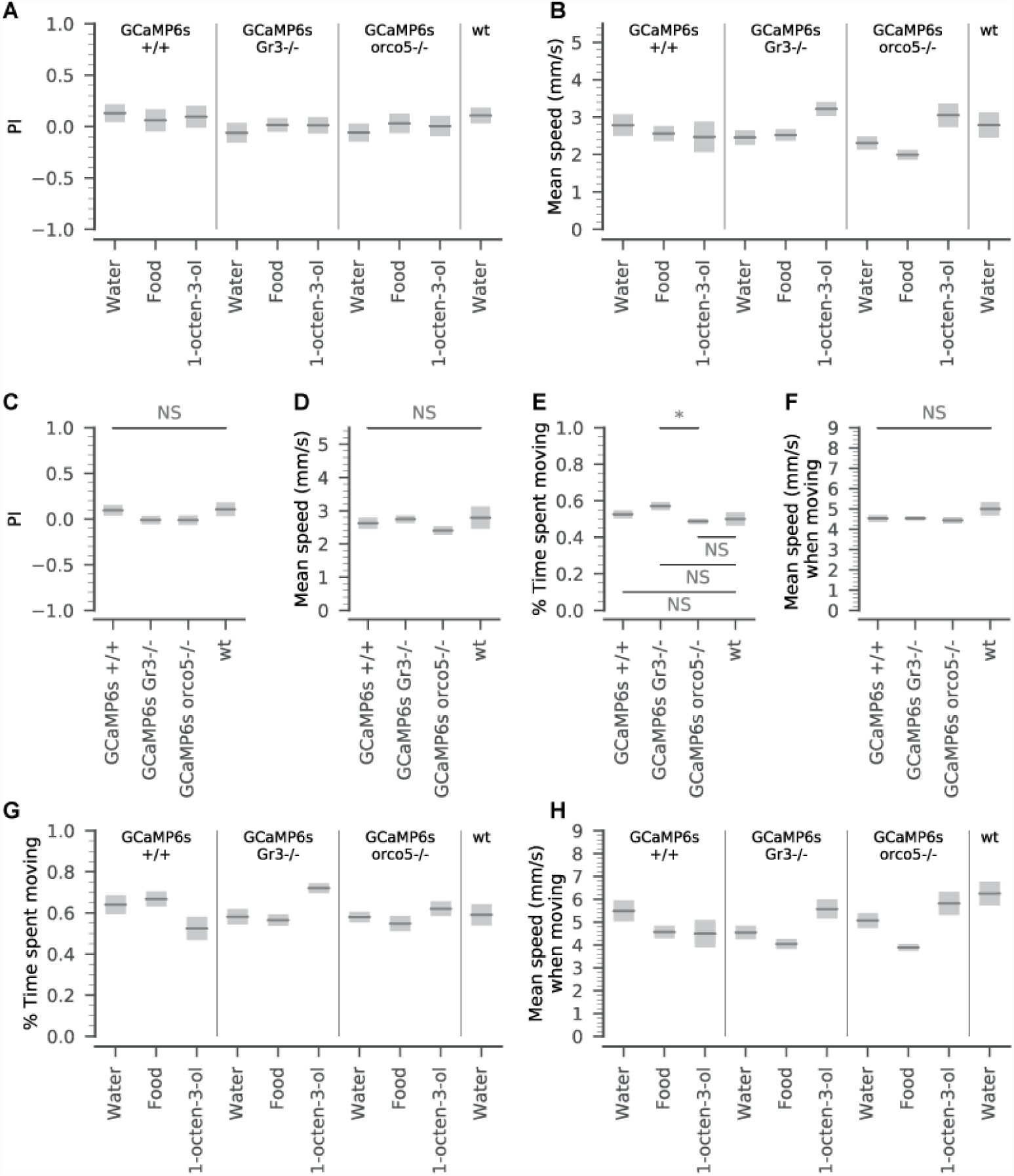
Innate behavioral kinematics in the acclimation phase of larval behavior assays. Prior to stimulation we found no differences in position (A) or mean speed (B) between larvae of the mutant backgrounds, and between the mutants and the Liverpool strain background (1-way ANOVA p-value = 0.315 [PI]; p-value = 0.3173 [mean speed]). In addition, we investigated additional parameters of larval movement before addition of experimental odors grouped by mutant strain. In the majority of variables (PI (C), mean speed (D)), we found no significant differences between strains (pairwise t-tests with Holm-Bonferroni correction). Although the proportion of time spent moving was significantly different between GCaMP6s/orco5-/- and GCaMP6s/Gr3-/- larvae ((E), p-value = 0.017), neither group was significantly different from the behaviors observed in wt Liverpool larvae. “Moving” was defined as translational movement > In the experiment phase, after the odor was introduced to the arena, the animals still did not exhibit significant differences in the proportion of time spent moving (G) (1-way ANOVA p-value = 0.478). However, their mean speed when moving was significantly different across strains and treatments (2-way ANOVA p-value = 0.007 background effect; p-value = 0.001 treatment effect; p-value = 0.054 interaction effect). When comparing the speed when moving > between food odor and the water control for each strain, GCaMP6s/orco5-/- exhibited a significantly lower speed (pairwise t-test p-value = 0.0037), and GCaMP6s+/+ mosquitos showed a trend toward a lower speed (pairwise t-test p-value = 0.092). However, GCaMP6s/Gr3-/- larvae did not show this effect (pairwise t-test p-value = 0.18).

**Supplementary Figure 6.**
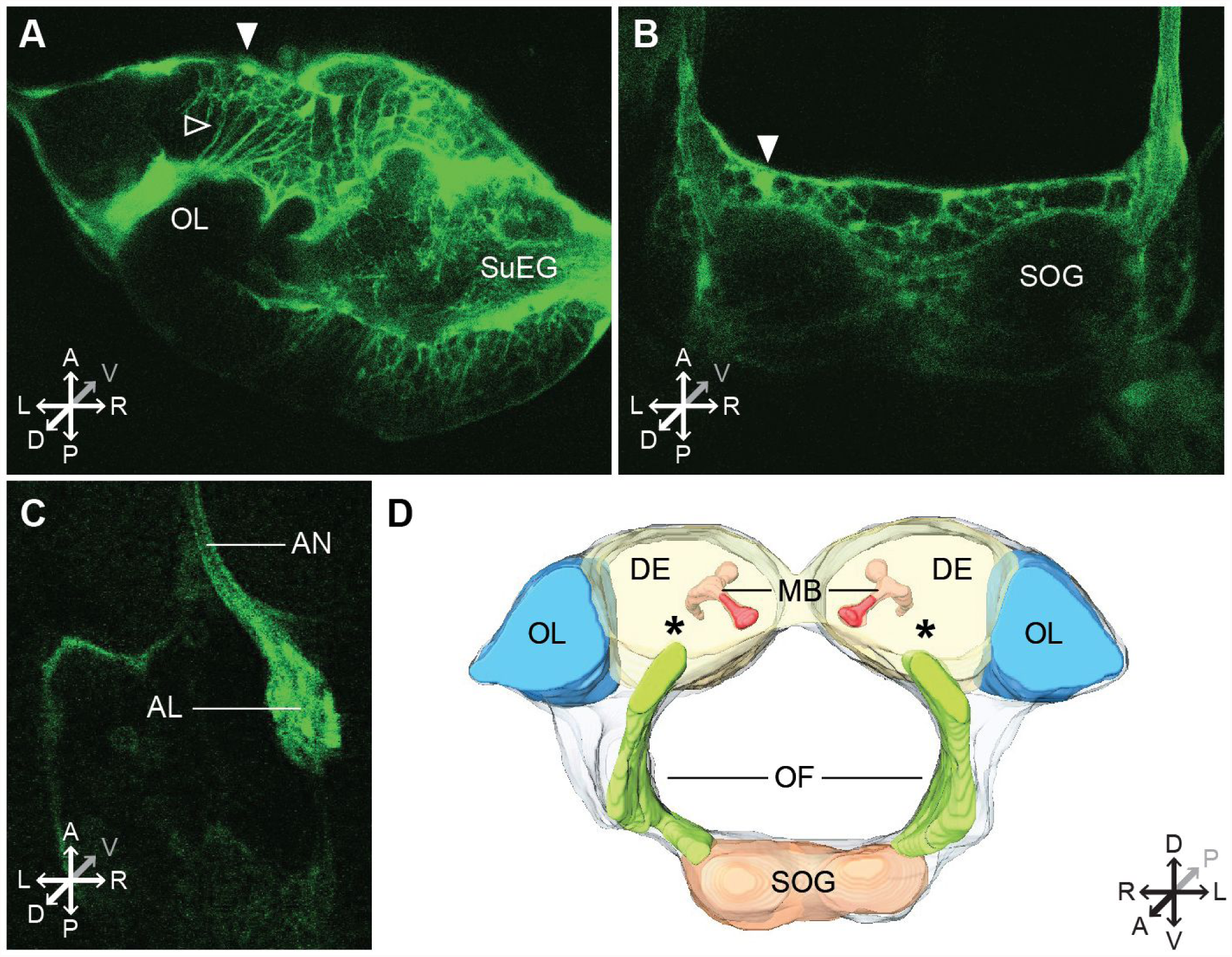
Two-photon imaging of *Ae. aegypti* larvae. To further explore the potential functionality of our GCaMP6s mosquito line, we used 2-photon microscopy to investigate higher spatial resolution in imaging GCaMP6s expression. Briefly, a larva was transferred to a Peltier-cooled holder that allows for the head to be fixed to the stage using ultraviolet glue. GCaMP6s expression was imaged at 2 Hz using the Prairie Ultima IV two-photon excitation microscope (Prairie Technologies) and Ti-Sapphire laser (Chameleon Ultra; Coherent). Although this experimental protocol was unsuitable for imaging stimulus-evoked responses due to the substantial movement of the brain, we were able to image various regions of interest throughout the brain of live, head-fixed larvae, including cell bodies (labeled with a filled arrow) and neuropil (labeled with an open arrow). These areas of interest included the optic lobe; OL, supra-esophageal ganglion; SuEG (upper DE, (A)), subesophageal ganglion; SOG (B), antennal nerve; AN and antennal lobe; AL (C). [A; B: L4 larvae with dorsal head cuticle removed. C: L2 larva imaged through transparent cuticle]. D: Approximate 3D reconstruction of larval brain regions based on a confocal scan of the dissected larval brain. Asterisks indicate the rough location of the antennal lobe. Additional labeled brain regions include a general schematic of the optic lobe (OL), deutocerebrum (DE), mushroom bodies (MB), suboesophageal ganglion (SOG), and oesophageous foramen (OF) [84],Thermo Scientific Amira Software).

**Supplementary Figure 7.**
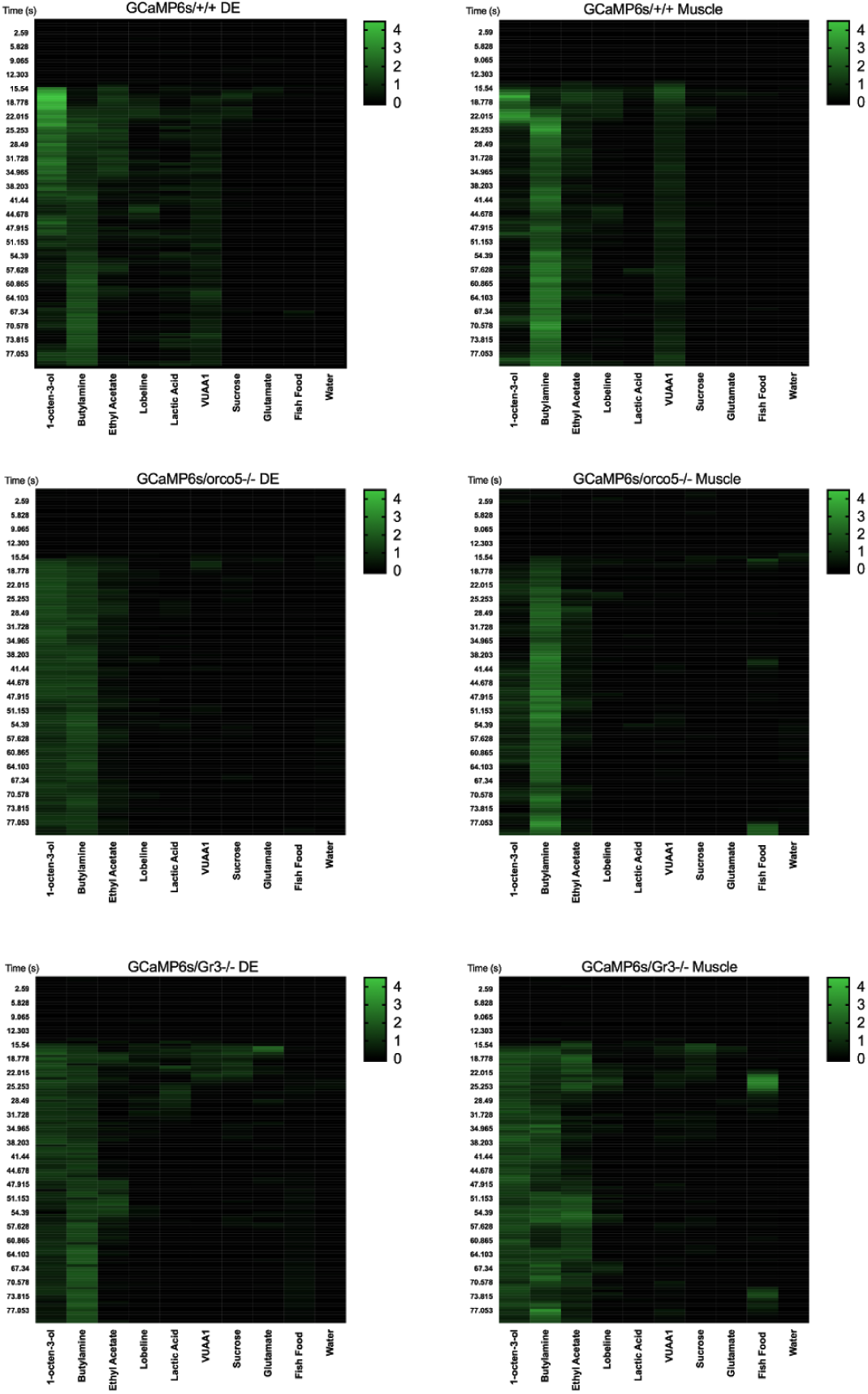
Average calcium responses of GCaMP6s+/+, GCaMP/orco5-/-, and GCaMP6s/Gr3-/- over time. Responses to various stimuli were averaged over multiple replicates for each time point.

**Supplementary Data 1.** The Python code used to analyze and interpret larval behavior assays, as well as raw trajectory data for each larval behavior experiment (Figure 3, Supplemental Figure 5) are available at https://github.com/eleanorlutz/aedes-aegypti-gcamp6s-larval-behavior

**Supplemental Video 1.** Time-lapse of female GCaMP6s/+/+ adult mouthparts taken on a Leica SP5 at 10x magnification.

**Supplemental Video 2.** Time-lapse of female GCaMP6s/+/+ adult antennae taken on a Leica SP5 at 10x magnification.

**Supplemental Video 3.** Time-lapse of an adult female GCaMP6s/+/+ right compound eye on a Leica SP5 at 10x magnification.

**Supplemental Video 4.** Time-lapse of a whole L2 GCaMP6s/+/+ larvae taken on a Leica SP5 at 5x magnification.

**Supplemental Video 5.** Time-lapse of a L2 GCaMP6s/+/+ larva head taken on a Leica SP5 at 10x magnification.

**Supplementary Table 1.**
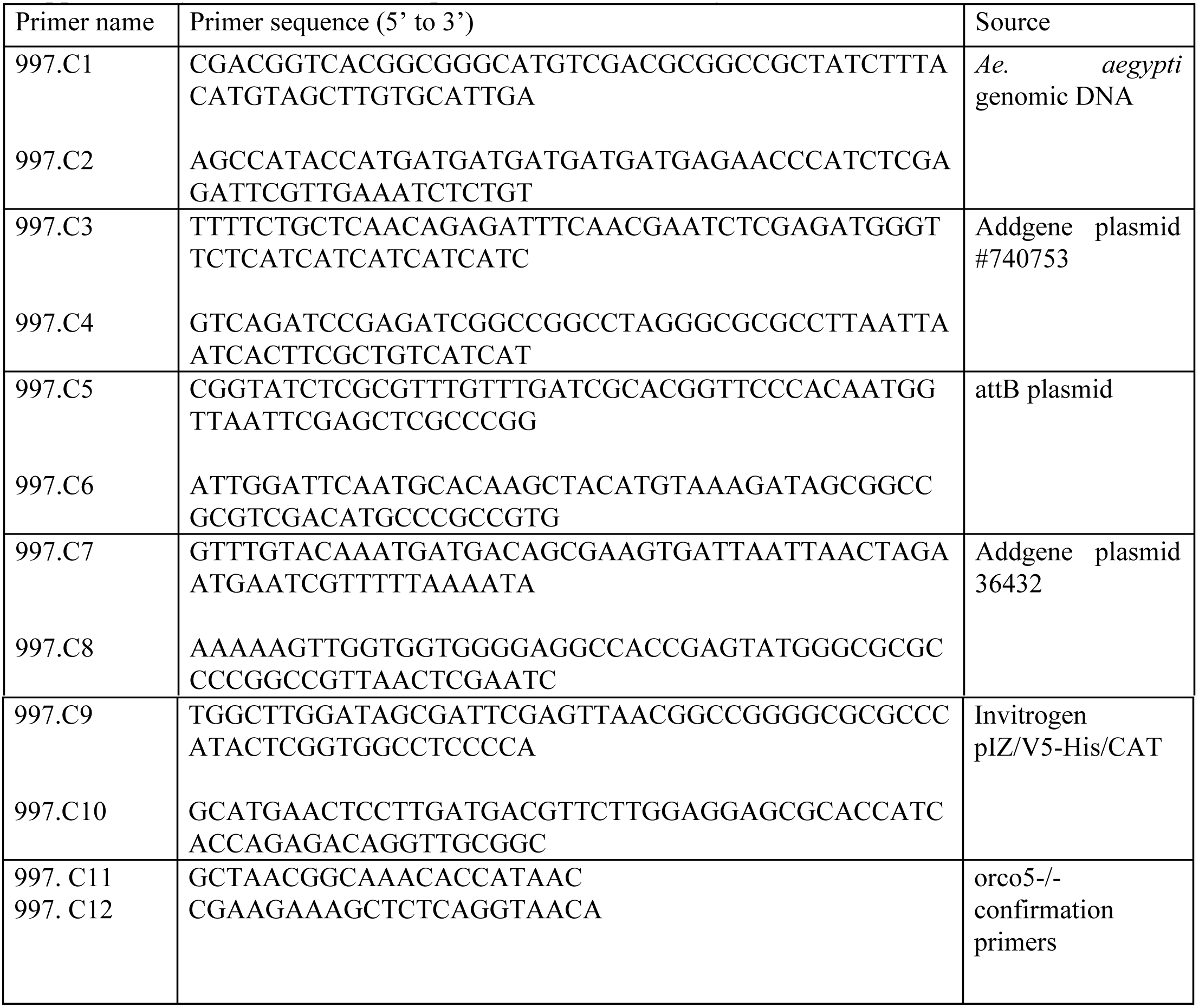
Primer Sequences U in this

**Supplementary Table 2.**
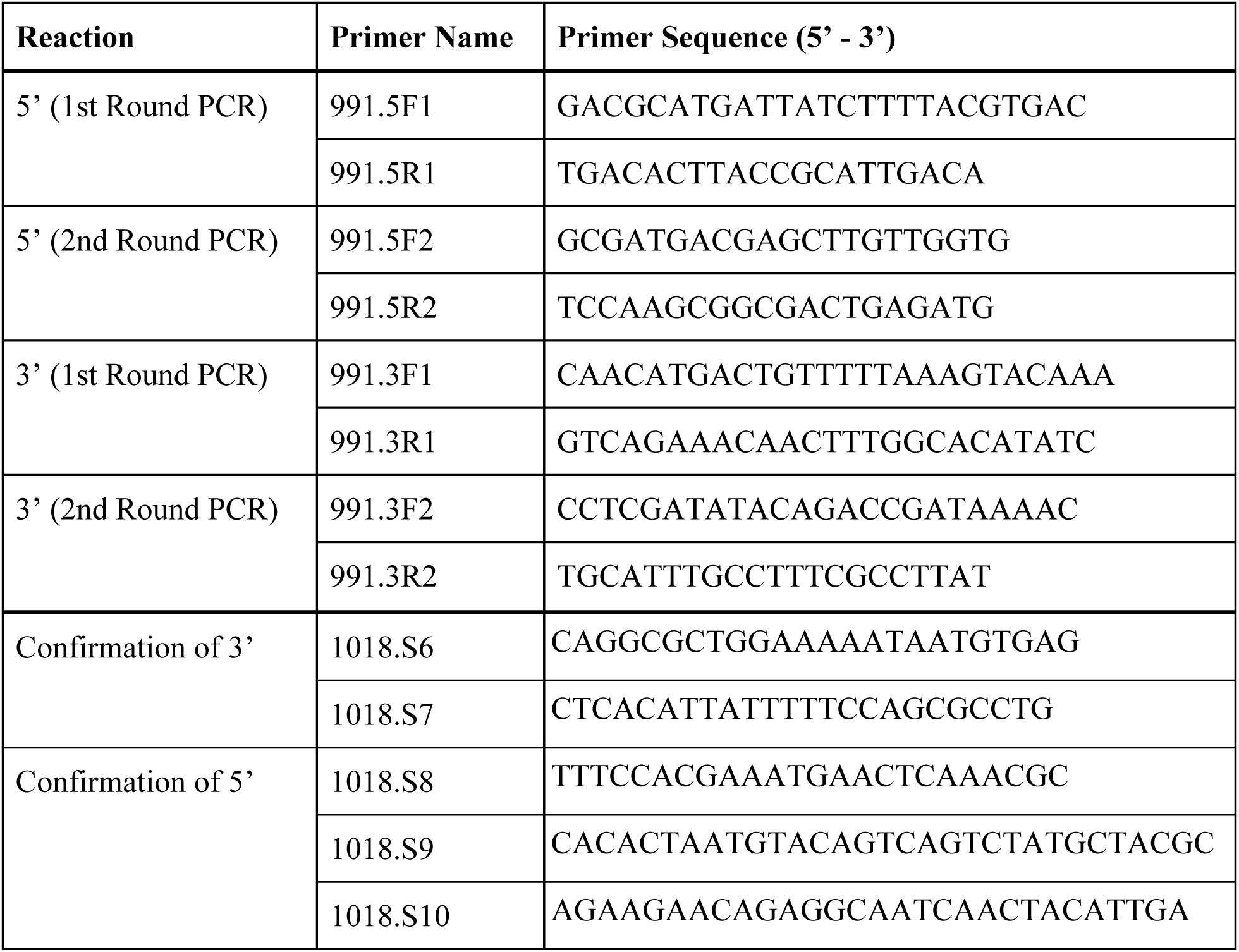
Inverse PCR Primer sequences used in this study

## References

1. Weaver SC, Charlier C, Vasilakis N, Lecuit M. Zika, Chikungunya, and Other Emerging Vector-Borne Viral Diseases. Annu Rev Med [Internet]. 2017; Available from: http://dx.doi.org/10.1146/annurev-med-050715-105122.

2. Barrett ADT, Higgs S. Yellow fever: a disease that has yet to be conquered. Annu Rev Entomol. 2007;52:209–29.

3. Halstead SB. Dengue virus-mosquito interactions. Annu Rev Entomol. 2008;53:273–91.

4. Weaver SC, Barrett ADT. Transmission cycles, host range, evolution and emergence of arboviral disease. Nat Rev Microbiol. 2004;2:789–801

5. Weaver SC, Reisen WK. Present and future arboviral threats. Antiviral Res. 2010;85:328–45.

6. Harris AF, McKemey AR, Nimmo D, Curtis Z, Black I, Morgan SA, et al. Successful suppression of a field mosquito population by sustained release of engineered male mosquitoes. Nat Biotechnol. 2012;30:828–30.

7. McMeniman CJ, Lane RV, Cass BN, Fong AWC, Sidhu M, Wang Y-F, et al. Stable introduction of a life-shortening Wolbachia infection into the mosquito Aedes aegypti. Science. 2009;323:141–4.

8. Walker T, Johnson PH, Moreira LA, Iturbe-Ormaetxe I, Frentiu FD, McMeniman CJ, et al. The wMel Wolbachia strain blocks dengue and invades caged Aedes aegypti populations. Nature. 2011;476:450–3.

9. Sinkins SP. Wolbachia and cytoplasmic incompatibility in mosquitoes. Insect Biochem Mol Biol. 2004;34:723–9.

10. Champer J, Buchman A, Akbari OS. Cheating evolution: engineering gene drives to manipulate the fate of wild populations. Nat Rev Genet. 2016;17:146–59.

11. Marshall JM, Akbari OS. Can CRISPR-Based Gene Drive Be Confined in the Wild? A Question for Molecular and Population Biology. ACS Chem Biol. 2018;13:424–30.

12. Marshall JM, Akbari OS. Gene Drive Strategies for Population Replacement. Genetic Control of Malaria and Dengue. 2016. p. 169–200.

13. Maoz D, Ward T, Samuel M, Müller P, Runge-Ranzinger S, Toledo J, et al. Community effectiveness of pyriproxyfen as a dengue vector control method: A systematic review. PLoS Negl Trop Dis. 2017;11:e0005651.

14. Moyes CL, Vontas J, Martins AJ, Ng LC, Koou SY, Dusfour I, et al. Contemporary status of insecticide resistance in the major Aedes vectors of arboviruses infecting humans. PLoS Negl Trop Dis. 2017;11:e0005625.

15. Franz AWE, Clem RJ, Passarelli AL. Novel Genetic and Molecular Tools for the Investigation and Control of Dengue Virus Transmission by Mosquitoes. Curr Trop Med Rep. 2014;1:21–31.

16. Montell C, Zwiebel LJ. Chapter Ten - Mosquito Sensory Systems. In: Raikhel AS, editor. Advances in Insect Physiology. Academic Press; 2016. p. 293–328.

17. Lutz EK, Lahondère C, Vinauger C, Riffell JA Olfactory learning and chemical ecology of olfaction in disease vector mosquitoes: a life history perspective. Curr Opin Insect Sci. 2017;20:75–83.

18. Syed Z. Chemical ecology and olfaction in arthropod vectors of diseases. Current Opinion in Insect Science. Elsevier; 2015;10:83–9.

19. DeGennaro M, McBride CS, Seeholzer L, Nakagawa T, Dennis EJ, Goldman C, et al. orco mutant mosquitoes lose strong preference for humans and are not repelled by volatile DEET. Nature. 2013;498:487–91.

20. McMeniman CJ, Corfas RA, Matthews BJ, Ritchie SA, Vosshall LB. Multimodal integration of carbon dioxide and other sensory cues drives mosquito attraction to humans. Cell. 2014;156:1060–71.

21. van Breugel F, Riffell J, Fairhall A, Dickinson MH. Mosquitoes Use Vision to Associate Odor Plumes with Thermal Targets. Curr Biol. 2015;25:2123–9.

22. Kennedy JS. The Visual Responses of Flying Mosquitoes. Proc Zool Soc Lond. 2009;A109:221–42.

23. Vinauger C, Lahondère C, Wolff GH, Locke LT, Liaw JE, Parrish JZ, et al. Modulation of Host Learning in Aedes aegypti Mosquitoes. Curr Biol. 2018;28:333–44e8.

24. Xia Y, Wang G, Buscariollo D, Pitts RJ, Wenger H, Zwiebel LJ. The molecular and cellular basis of olfactory-driven behavior in Anopheles gambiae larvae. Proc Natl Acad Sci U S A. 2008;105:6433–8.

25. Liu C, Jason Pitts R, Bohbot JD, Jones PL, Wang G, Zwiebel LJ. Distinct Olfactory Signaling Mechanisms in the Malaria Vector Mosquito Anopheles gambiae. PLoS Biol. 2010;8:e1000467.

26. Christophers S. Aedes aegypti (L.) the yellow fever mosquito: its life history, bionomics and structure. aegypti (L) the Yellow Fever Mosquito: its Life…Internet]. cabdirect.org;1960; Available from: https://www.cabdirect.org/cabdirect/abstract/19602901825.

27. Ghaninia M, Larsson M, Hansson BS, Ignell R. Natural odor ligands for olfactory receptor neurons of the female mosquito Aedes aegypti: use of gas chromatography-linked single sensillum recordings. J Exp Biol. 2008;211:3020–7.

28. Moffett DF, Jagadeshwaran U, Wang Z, Davis HM, Onken H, Goss GG. Signaling by intracellular Ca2 and H in larval mosquito (Aedes aegypti) midgut epithelium in response to serosal serotonin and lumen pH. J Insect Physiol. 2012;58:506–12.

29. Macleod GT, Hegström-Wojtowicz M, Charlton MP, Atwood HL. Fast calcium signals in Drosophila motor neuron terminals. J Neurophysiol. 2002;88:2659–63.

30. Chen T-W, Wardill TJ, Sun Y, Pulver SR, Renninger SL, Baohan A, et al. Ultrasensitive fluorescent proteins for imaging neuronal activity. Nature. 2013;499:295–300.

31. Anderson MAE, Gross TL, Myles KM, Adelman ZN. Validation of novel promoter sequences derived from two endogenous ubiquitin genes in transgenic Aedes aegypti. Insect Mol Biol. 2010;19:441–9.

32. Akbari OS, Antoshechkin I, Amrhein H, Williams B, Diloreto R, Sandler J, et al. The developmental transcriptome of the mosquito Aedes aegypti, an invasive species and major arbovirus vector. G3. 2013;3:1493–509.

33. Taylor RW, Romaine IM, Liu C, Murthi P, Jones PL, Waterson AG, et al. Structure–Activity Relationship of a Broad-Spectrum Insect Odorant Receptor Agonist. ACS Chem Biol. 2012;7:1647–52.

34. Reddy GVP, Guerrero A. Interactions of insect pheromones and plant semiochemicals. Trends Plant Sci. 2004;9:253–61.

35. Acree F, Turner RB, Gouck HK, Beroza M, Smith N. L-Lactic Acid: A Mosquito Attractant Isolated from Humans. Science. 1968;161:1346–7.

36. Hoel DF, Kline DL, Allan SA, Grant A. Evaluation of carbon dioxide, 1-octen-3-ol, and lactic acid as baits in Mosquito Magnet Pro traps for Aedes albopictus in north central Florida. J Am Mosq Control Assoc. 2007;23:11–7.

37. Gonzalez PV, González Audino PA, Masuh HM. Behavioral Response of Aedes aegypti (Diptera: Culicidae) Larvae to Synthetic and Natural Attractants and Repellents. J Med Entomol. 2015;52:1315–21.

38. Freeman EG, Wisotsky Z, Dahanukar A. Detection of sweet tastants by a conserved group of insect gustatory receptors. Proc Natl Acad Sci U S A. 2014;111:1598–603.

39. Montell C. A taste of the Drosophila gustatory receptors. Curr Opin Neurobiol. 2009;19:345–53.

40. Chitarra GS, Abee T, Rombouts FM, Posthumus MA, Dijksterhuis J. Germination of penicillium paneum Conidia is regulated by 1-octen-3-ol, a volatile self-inhibitor. Appl Environ Microbiol. 2004;70:2823–9.

41. Broek IVF, Otter CJ. Odour sensitivity of antennal olfactory cells underlying grooved pegs of Anopheles gambiae s.s. and An. quadriannulatus. Entomol Exp Appl. 2000;96:167–75.

42. Kumar BN, Taylor RW, Pask GM, Zwiebel LJ, Newcomb RD, Christie DL. A conserved aspartic acid is important for agonist (VUAA1) and odorant/tuning receptor-dependent activation of the insect odorant co-receptor (Orco). PLoS One. 2013;8:e70218.

43. Bohbot JD, Dickens JC. Characterization of an enantioselective odorant receptor in the yellow fever mosquito Aedes aegypti. PLoS One. 2009;4:e7032.

44. Erdelyan CNG, Mahood TH, Bader TSY, Whyard S. Functional validation of the carbon dioxide receptor genes in Aedes aegypti mosquitoes using RNA interference. Insect Mol Biol. 2012;21:119–27.

45. Bohbot J, Pitts RJ, Kwon H-W, Rützler M, Robertson HM, Zwiebel LJ. Molecular characterization of the Aedes aegypti odorant receptor gene family. Insect Mol Biol. Wiley Online Library; 2007;16:525–37.

46. Xia Y, Wang G, Buscariollo D, Pitts RJ, Wenger H, Zwiebel LJ. The molecular and cellular basis of olfactory-driven behavior in Anopheles gambiae larvae. Proc Natl Acad Sci U S A. National Acad Sciences; 2008;105:6433–8.

47. Gonzalez PV, González Audino PA, Masuh HM. Behavioral Response of Aedes aegypti (Diptera: Culicidae) Larvae to Synthetic and Natural Attractants and Repellents. J Med Entomol. 2015;52:1315–21.

48. Merritt RW, Dadd RH, Walker ED. Feeding behavior, natural food, and nutritional relationships of larval mosquitoes. Annu Rev Entomol. 1992;37:349–76.

49. Riffell JA, Abrell L, Hildebrand JG. Physical processes and real-time chemical measurement of the insect olfactory environment. J Chem Ecol. 2008;34:837–53.

50. Burks RL, Lodge DM. Cued in: advances and opportunities in freshwater chemical ecology. J Chem Ecol. 2002;28:1901–17.

51. Mysore K, Flannery EM, Tomchaney M, Severson DW, Duman-Scheel M. Disruption of Aedes aegypti olfactory system development through chitosan/siRNA nanoparticle targeting of semaphorin-1a. PLoS Negl Trop Dis. journals.plos.org; 2013;7e2215.

52. Mysore K, Flister S, Müller P, Rodrigues V, Reichert H. Brain development in the yellow fever mosquito Aedes aegypti: a comparative immunocytochemical analysis using cross-reacting antibodies from Drosophila melanogaster. Dev Genes Evol. 2011;221:281–96.

53. Crespo JG. A review of chemosensation and related behavior in aquatic insects. J Insect Sci. 2011;11:62.

54. Mellon D Jr. Combining dissimilar senses: central processing of hydrodynamic and chemosensory inputs in aquatic crustaceans. Biol Bull. 2007;213:1–11

55. Thiel M, Breithaupt T. Chemical Communication in Crustaceans: Research Challenges for the Twenty-First Century. In: Breithaupt T.T, Thiel M, editors. Chemical Communication in Crustaceans. New York, NY: Springer New York;2011.p.3–22.

56. Takken W, Kline DL. Carbon dioxide and 1-octen-3-ol as mosquito attractants. J Am Mosq Control Assoc. 1989;5:311–6.

57. Fischer G, Dott W. Relevance of airborne fungi and their secondary metabolites for environmental, occupational and indoor hygiene. Arch Microbiol. 2003;179:75–82.

58. Gomez-Marin A, Stephens GJ, Louis M. Active sampling and decision making in Drosophila chemotaxis. Nat Commun. 2011;2:441.

59. Sourjik V, Wingreen NS. Responding to chemical gradients: bacterial chemotaxis. Curr Opin Cell Biol. 2012;24:262–8.

60. Hilliard MA, Bargmann CI, Bazzicalupo P. C. elegans responds to chemical repellents by integrating sensory inputs from the head and the tail. Curr Biol. 2002;12:730–4.

61. McMeniman CJ, Corfas RA, Matthews BJ, Ritchie SA, Vosshall LB. Multimodal integration of carbon dioxide and other sensory cues drives mosquito attraction to humans. Cell. 2014;156:1060–71.

62. van Breugel F, Riffell J, Fairhall A, Dickinson MH. Mosquitoes Use Vision to Associate Odor Plumes with Thermal Targets. Curr Biol. 2015;25:2123–9.

63. Bräcker LB, Siju KP, Varela N. Aso Y, Zhang M, Hein I, et al. Essential Role of the Mushroom Body in Context-Dependent CO2 Avoidance in Drosophila. Curr Biol. 2013;23:1228–34.

64. Lewis LPC, Siju KP, Aso Y, Friedrich AB, Bulteel AJB, Rubin GM, et al. A Higher Brain Circuit for Immediate Integration of Conflicting Sensory Information in Drosophila. Curr Biol. 2015;25:2203–14.

65. Riabinina O, Task D, Marr E, Lin C-C, Alford R, O’Brochta DA, et al. Organization of olfactory centres in the malaria mosquito Anopheles gambiae. Nat Commun. nature.com; 2016;7:13010.

66. Chen C-L, Hermans L, Viswanathan MC, Fortun D, Unser M, Cammarato A, et al. Imaging neural activity in the ventral nerve cord of behaving adult Drosophila [Internet]. bioRxiv. 2018 [cited 2018 May 4]. p. 250118. Available from: https://www.biorxiv.org/content/early/2018/01/22/250118.abstract

67. Schnell B, Ros IG, Dickinson MH. A Descending Neuron Correlated with the Rapid Steering Maneuvers of Flying Drosophila. Curr Biol. 2017;27:1200–5.

68. Kim AJ, Fenk LM, Lyu C, Maimon G. Quantitative Predictions Orchestrate Visual Signaling in Drosophila. Cell. 2017;168:280–94e12.

69. Pelletier J, Guidolin A, Syed Z, Cornel AJ, Leal WS. Knockdown of a mosquito odorant-binding protein involved in the sensitive detection of oviposition attractants. J Chem Ecol. 2010;36:245–8.

70. Li M, Bui M, Yang T, Bowman CS, White BJ, Akbari OS. Germline Cas9 expression yields highly efficient genome engineering in a major worldwide disease vector, Aedes aegypti. Proc Natl Acad Sci U S A. 2017;114:E10540–9.

71. Gibson DG, Young L, Chuang R-Y, Craig Venter J, Hutchison CA, Smith HO. Enzymatic assembly of DNA molecules up to several hundred kilobases. Nat Methods. Nature Publishing Group; 2009;6:343–5.

72. Bischof J, Maeda RK, Hediger M, Karch F, Basler K. An optimized transgenesis system for Drosophila using germ-line-specific ϕC31 integrases. Proceedings of the National Academy of Sciences. 2007;104:3312–7.

73. Anderson MAE, Gross TL, Myles KM, Adelman ZN. Validation of novel promoter sequences derived from two endogenous ubiquitin genes in transgenic Aedes aegypti. Insect Mol Biol. 2010;19:441–9.

74. Pfeiffer BD, Truman JW, Rubin GM. Using translational enhancers to increase transgene expression in Drosophila. Proc Natl Acad Sci U S A. 2012;109:6626–31.

75. Theilmann DA, Stewart S. Tandemly repeated sequence at the 3’ end of the IE-2 gene of the baculovirus Orgyia pseudotsugata multicapsid nuclear polyhedrosis virus is an enhancer element. Virology. 1992;187:97–106.

76. Handler AM, Harrell RA Ii. Germline transformation of Drosophila melanogaster with the piggyBac transposon vector. Insect Mol Biol. 1999;8:449–57.

77. Kokoza V, Ahmed A, Wimmer EA, Raikhel AS. Efficient transformation of the yellow fever mosquito Aedes aegypti using the piggyBac transposable element vector pBac[3xP3-EGFP afm]. Insect Biochem Mol Biol. 2001;31:1137–43.

78. Lobo NF, Hua-Van A, Li X, Nolen BM, Fraser MJ Jr. Germ line transformation of the yellow fever mosquito, Aedes aegypti, mediated by transpositional insertion of a piggyBac vector. Insect Mol Biol. 2002;11:133–9.

79. Akbari OS, Papathanos PA, Sandler JE, Kennedy K, Hay BA. Identification of germline transcriptional regulatory elements in Aedes aegypti. Sci Rep. 2014;4:3954

80. Aryan A, Myles KM, Adelman ZN. Targeted genome editing in Aedes aegypti using TALENs. Methods. 2014;69:38–45.

81. Huang AM, Rehm EJ, Rubin GM. Recovery of DNA sequences flanking P-element insertions in Drosophila: inverse PCR and plasmid rescue. Cold Spring Harb Protoc. 2009;2009:db.prot5199.

82. Jia H, Rochefort NL, Chen X, Konnerth A. In vivo two-photon imaging of sensory-evoked dendritic calcium signals in cortical neurons. Nat Protoc. 2011;6:28–35.

83. Schindelin J, Arganda-Carreras I, Frise E, Kaynig V, Longair M, Pietzsch T, et al. Fiji: an open-source platform for biological-image analysis. Nat Methods. 2012;9:676–82.

84. Mysore K, Flister S, Müller P, Rodrigues V, Reichert H. Brain development in the yellow fever mosquito Aedes aegypti: a comparative immunocytochemical analysis using cross-reacting antibodies from Drosophila melanogaster. Dev Genes Evol. 2011;221:281–96.

